# Library-based single-cell analysis of CAR signaling reveals drivers of *in vivo* persistence

**DOI:** 10.1101/2024.04.29.591541

**Authors:** Caleb R. Perez, Andrea Garmilla, Avlant Nilsson, Hratch M. Baghdassarian, Khloe S. Gordon, Louise G. Lima, Blake E. Smith, Marcela V. Maus, Douglas A. Lauffenburger, Michael E. Birnbaum

**Affiliations:** Koch Institute for Integrative Cancer Research, Cambridge, MA, USA; Department of Biological Engineering, Massachusetts Institute of Technology, Cambridge, MA, USA; Singapore-MIT Alliance for Research and Technology Centre, Singapore, Singapore; Program in Immunology, Harvard Medical School, Boston, MA, USA; Department of Biology and Biological Engineering, Chalmers University of Technology, Gothenburg, SE, Sweden; Cellular Immunotherapy Program, Massachusetts General Hospital Cancer Center, Boston, MA, USA; Department of Medicine, Harvard Medical School, Boston, MA, USA; Ragon Institute of MIT, MGH and Harvard, Cambridge, MA, USA

**Keywords:** T cells, T cell signaling, immunotherapy, CAR T cells, chimeric antigen receptors, intracellular signaling domains, pooled screens, single-cell RNA sequencing, tonic signaling, persistence

## Abstract

The anti-tumor function of engineered T cells expressing chimeric antigen receptors (CARs) is dependent on signals transduced through intracellular signaling domains (ICDs). Different ICDs are known to drive distinct phenotypes, but systematic investigations into how ICD architectures direct T cell function—particularly at the molecular level—are lacking. Here, we use single-cell sequencing to map diverse signaling inputs to transcriptional outputs, focusing on a defined library of clinically relevant ICD architectures. Informed by these observations, we functionally characterize transcriptionally distinct ICD variants across various contexts to build comprehensive maps from ICD composition to phenotypic output. We identify a unique tonic signaling signature associated with a subset of ICD architectures that drives durable *in vivo* persistence and efficacy in liquid, but not solid, tumors. Our findings work toward decoding CAR signaling design principles, with implications for the rational design of next-generation ICD architectures optimized for *in vivo* function.

## Introduction

The engineering of T cells to express chimeric antigen receptors (CARs) and other synthetic signaling constructs has transformed cancer treatment. By directing the natural trafficking, immunostimulatory, and cytotoxic functions of T cells toward specific tumor antigens, CARs enable potent anti-tumor responses.^1^ This has been most successful in relapsed and refractory B cell malignancies, driving complete and durable remissions in the majority of patients.^2,3^ However, these therapies are marred by significant toxicities and frequent relapse,^4,5^ and the successes observed in liquid cancers have yet to be meaningfully translated to solid tumors.^6^ Given these challenges, optimizing the function and phenotype of CAR T cells remains an important clinical need.

CARs mediate tumor-specific responses by linking a tumor-binding moiety—most commonly a single chain variable fragment (scFv) against CD19 or BCMA in B cell cancers—to immunostimulatory intracellular signaling domains (ICDs), driving signaling cascades upon antigen binding that direct T cell activation, proliferation, and effector function.^7^ The specific identity and arrangement of these ICDs has been linked to varying phenotypic effects across numerous clinical and preclinical studies. All approved CAR products utilize one of two combinations of ICDs, with the CD3z ICD providing an activating signal derived from the T cell receptor (TCR) complex, and either a CD28 or 4-1BB ICD providing costimulation (referred to as 28z or BBz, respectively).^3^ Although both have demonstrated strong efficacy profiles in patients, 28z CAR T cells are thought to exhibit quicker expansion kinetics, greater peak expansion, but reduced persistence relative to BBz CARs,^2,7^ functional characteristics attributed to differences in signaling pathways, potency, and kinetics that favor an effector memory over central memory state.^8–10^ These results highlight the impact of even relatively minor changes in ICDs.

Outside of these well-studied ICD compositions, a growing number of studies have shown that incorporating additional or alternative ICDs into CARs induces phenotypes distinct from either 28z or BBz. These include incorporating alternative T cell signals, such as CD3e, and costimulatory ICDs beyond CD28 and 4-1BB;^11–14^ making rational mutations to key signaling motifs, such as inactivating immunoreceptor tyrosine-based activation motifs (ITAMs) in CD3z to finetune signaling;^15,16^ adding cytokine receptor-derived signals to provide antigen-mediated induction of JAK-STAT signaling;^17,18^ and incorporating signals not typically expressed by T cells to access non-endogenous signaling programs, including those of myeloid,^19,20^ NK cell,^21–24^ and B cell origin.^19,25^ Collectively, these engineering strategies have reduced exhaustion, boosted persistence, enhanced antigen sensitivity, and, in some cases, improved preclinical efficacy.

Although these reports link ICD architecture to T cell phenotype and function, it remains unclear how variability in signaling inputs directs these responses at the molecular level. These studies have typically involved one-to-one comparisons to BBz and/or 28z benchmarks, and were tested across a wide range of cancer antigens, target binding affinities, tumor models, and CAR manufacturing protocols, limiting direct comparison across studies that could produce implementable design principles for CAR-T therapies. Recent work, including from our group, has probed randomized libraries of DNA-barcoded ICD compositions to select for defined characteristics like activation capacity, persistence, or killing,^26–28^ but these selections generally rely on a small subset of surface markers and thus do not provide global insight into the molecular drivers of these distinct phenotypes. Another report extended randomized ICD libraries to a single-cell RNA sequencing (scRNA-seq) readout, allowing deeper analysis of ICD-intrinsic transcriptional responses;^29^ however, phenotypic characterization was limited, particularly in terms of *in vivo* efficacy, making it difficult to assess functional consequences. We posit that a more thorough mapping of ICD input to functional output could reveal key ICD-intrinsic molecular drivers of CAR T cell function and efficacy, which could in turn accelerate the rational design of optimized CAR signaling.

To this end, here we report the design and validation of **caR**NA-seq, a library-based scRNA-seq platform for the high-throughput characterization of different CAR signaling inputs. We apply this platform to build a series of robust molecular datasets describing the transcriptional programs induced by a defined library of clinically relevant ICD architectures. By combining this data with thorough functional and phenotypic measurements on a subset of the library, we identify a unique tonic signaling signature, likely mediated by the CD40 ICD in a context-dependent manner, associated with long-term *in vivo* persistence and liquid tumor control. In addition, we use caRNA-seq to directly characterize disruptions in ICD-intrinsic transcriptional responses specific to the *in vivo* solid tumor microenvironment, which argues for the engineering of ICD architectures specifically for solid tumors. Together, these findings have broad implications for the design of next-generation CAR signaling architectures, both in liquid and solid tumors.

## Results

### Establishing a single-cell omics pipeline for the simultaneous characterization of diverse libraries of CAR signals

To develop a scRNA-seq-based pipeline for the rapid characterization of diverse CAR signaling inputs, we first sought to design a library of ICD architectures where CAR identity and CAR-T transcriptome could be readily captured in a single-cell workflow (**Figure 1A-C**). To maximize our ability to contextualize previous work in the development and characterization of novel CAR architectures, we built a library of ICD compositions that have been previously reported in preclinical and clinical studies, with a focus on those reported to drive distinct phenotypic effects. We created 36 distinct constructs (**Figure 1B** and **Table S1**), including a non-signaling negative control without ICDs (CAR1), sampling a diverse range of ICDs from different cell types of origin and functional families (**Table S2**). This library design has several advantages in the context of this study: it samples diverse signaling space while maintaining a feasible size given the cell number limitations of scRNA-seq; it ensures the majority of the library should functionally signal; and it maximizes the likelihood of observing differential phenotype and function across the library. To ensure we could ascribe any signaling or phenotypic differences to ICD composition rather than to factors such as target identity or binding affinity, we standardized each set of ICDs to use a single CD19-targeted binding domain/hinge domain/transmembrane domain backbone. Of the 36 variants, 35 expressed on the surface of Jurkat cell lines (**Figure S1A**); CAR22, the single exception, was removed from the library for downstream experiments.

**Figure 1.**
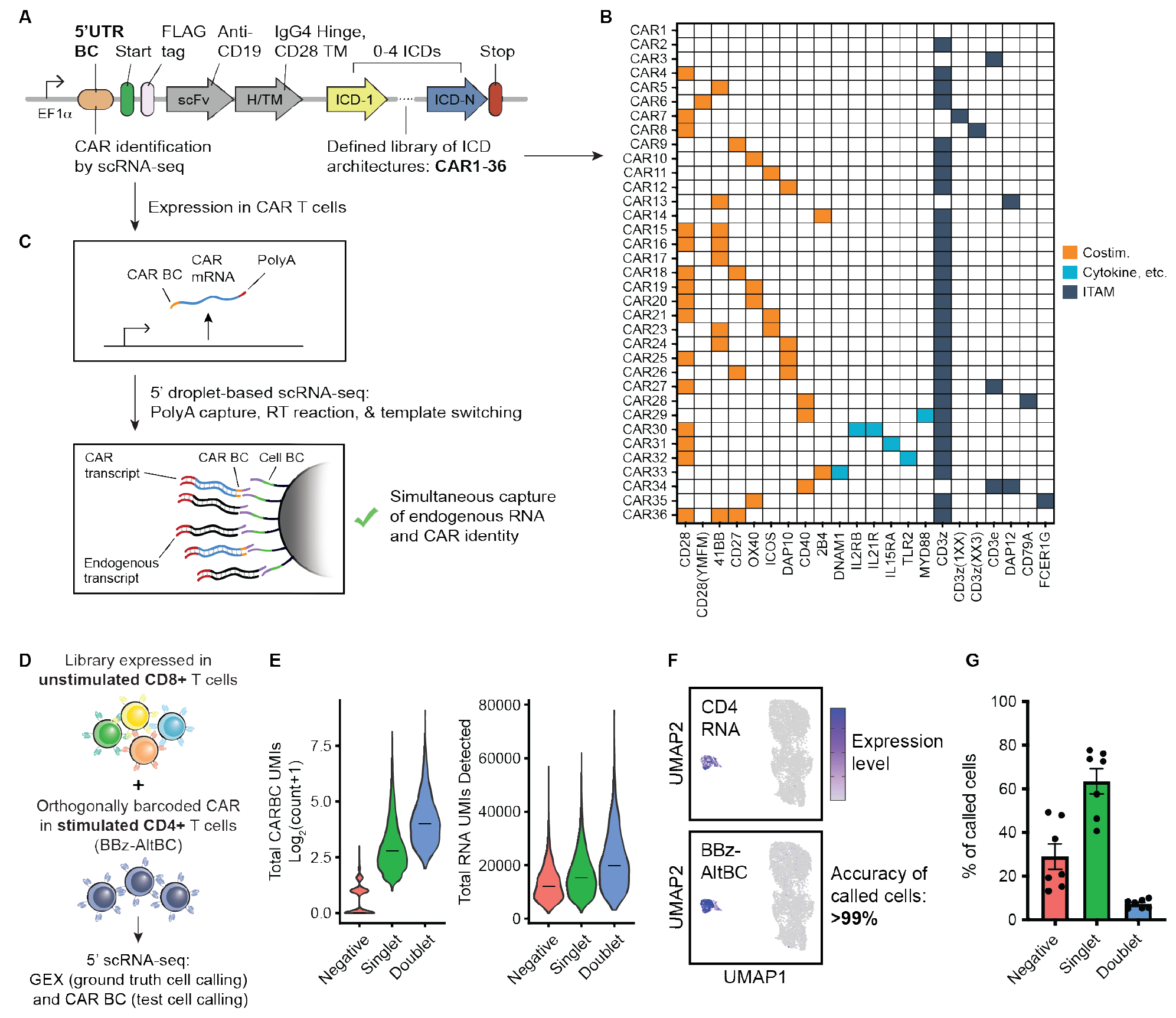
Design and validation of caRNA-seq, a high-throughput single-cell omics platform for the simultaneous transcriptional characterization of CAR signaling libraries. (A-C) Overview of the caRNA-seq platform. A defined library of CD19-targeted CARs of varied ICD architectures are labeled with barcodes in the 5’UTR, allowing simultaneous measurement of CAR identity and endogenous transcriptome within conventional 5’ capture droplet-based scRNA-seq workflows. CAR construct design is shown in (A), library design is shown in (B), and the barcoding schema is depicted in (C). (D) Design of CD4 spike-in experiment to assess accuracy of barcoding system. The caRNA- seq is expressed in unstimulated CD8 T cells, while an orthogonally barcoded BBz (BBz-AltBC) CAR is expressed in stimulated CD4 T cells, allowing assessment of the accuracy of CAR BC calling by comparison to ground truth identification of cell populations by gene expression (GEX). (E) Violin plots depicting CAR BC UMI counts and total RNA UMI counts of negative, singlet, and doublet populations following CAR calling algorithm. Bars represent median values across all cells within each group. (F) UMAP embeddings of CD4 and CD8 pools confidently called to a single CAR (n = 5,547 cells). Normalized RNA expression of CD4 and UMI counts of the alternate BBz CAR BC (BBz- AltBC) are overlaid. (G) Average recovery rates of single CARs across seven scRNA-seq experiments reported in this study, including negative and doublet rates. Data is presented as mean ± SEM.

To ensure each CAR variant could be linked to cell transcriptome, we designed a DNA barcoding schema capable of assigning ICD identity within conventional droplet-based scRNA-seq workflows (**Figure 1C**). Each CAR variant was modified to include a unique 8-bp barcode in the 5’UTR of its transgene. This CAR barcode (CAR BC) is expressed on the CAR transcript itself, which can be captured simultaneously alongside other polyadenylated transcripts. Its proximity to the transcription start site allows convenient sequencing of both the CAR BC and the cell barcode using commercially available 5’ capture kits (e.g. 10X Genomics) after simple PCR-based enrichment (see **Methods**), allowing measurement of single-cell gene expression and CAR identity within one assay. Using Jurkats, we validated that barcoding of the 5’UTR did not impact vector titer, CAR surface expression, or signaling (**Figure S1B-D**).

To assess the accuracy and sensitivity of the caRNA-seq system at library scale, we designed a controlled experiment in which gene expression served as ground truth: We expressed the full library in human CD8 T cells and an orthogonally barcoded 19BBz CAR (BBz-AltBC) in CD4 T cells, then we stimulated only the BBz-AltBC cells with CD19-expressing NALM6 leukemia cells. We then used 5’ scRNA-seq to sequence gene expression and CAR BCs of both CD4 and CD8 T cells in pool (**Figure 1D**). We recovered sufficient CAR BC UMIs to computationally call cells expressing a single CAR construct, while filtering out negatives and multiplets (**Figure 1E**; see **Methods**). To approximate the accuracy of CAR calling, we clustered all single CARs on the basis of gene expression, revealing a distinct activated CD4 T cell cluster, as expected (**Figure S1E- F**). BBz-AltBC CAR BC expression was almost entirely restricted to the CD4 cluster (**Figure 1F**), demonstrating the accuracy of our barcoding approach. In addition, within the unstimulated CD8 cluster, we were able to identify cells expressing every CAR variant in the library (**Figure S1G**), indicating that this approach is feasible at library scale. Supported by these initial results, we used the caRNA-seq platform to generate a series of transcriptional datasets to characterize our CAR ICD library under various conditions (**Table S3**). We observe a singlet CAR calling rate of >60% of cells across our datasets (**Figure 1G**), which is similar in performance to other direct polyadenylated transcript capture techniques.^30,31^

### Building a robust transcriptional dataset to map CAR signaling inputs to transcriptional output

To investigate the transcriptional landscape defined by the ICD architectures that comprise our library, we employed an *in vitro* model of repeated antigen rechallenge, which has been shown by our group and others to phenotypically differentiate CAR designs.^26,32^ T cells expressing each CAR variant were repeatedly exposed to NALM6 leukemia cells over a one-week period in arrayed cocultures, after which all variants were pooled equally and sequenced via caRNA-seq (**Figure 2A**). To avoid tumor cell outgrowth in the negative control culture, CAR1 cells were included after only a single NALM6 challenge on day 5. All other CARs controlled initial tumor challenges; however, a subset of the library began to lose tumor control by the end of the one-week coculture period (**Figure S2A**), indicating that this experimental paradigm successfully drives exhaustion and dysfunction for weaker library variants. To generate a robust dataset that fully captures ICD-intrinsic states, the library was challenged in array to eliminate crosstalk between different CARs. A comparison between the same batch of CARs challenged in array and in pool shows that the composition of cell states is meaningfully changed in the pooled context (**Figure S2B-D**), highlighting the importance of this approach.

**Figure 2.**
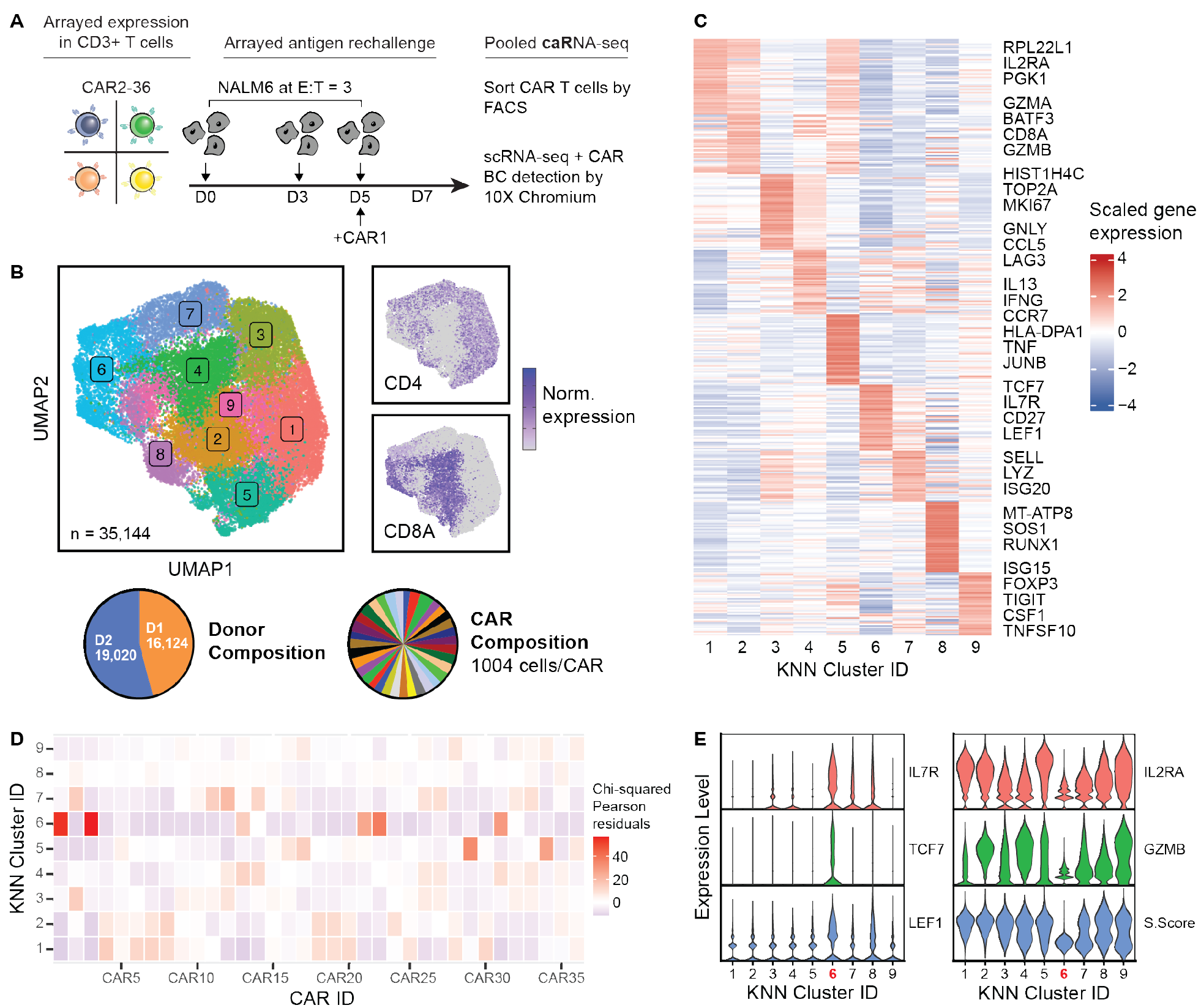
Building a robust *in vitro* dataset to link transcriptional profiles to CAR signaling inputs via caRNA-seq. (A) Experimental design for *in vitro* rechallenge assay. The caRNA-seq library was challenged in array with NALM6 cells three times over a seven-day period at an E:T ratio of 1:3. Cells were then pooled to sort for CARs, which were then input into scRNA-seq. CAR1 was subjected to only a single challenge at D5 to prevent tumor outgrowth in the negative control culture. (B) UMAP embeddings of all cells from two healthy donors, which were confidently called to a single CAR variant (n *=* 35,144 cells). Cells are colored by cell state (left), as determined by unsupervised clustering (K-Nearest Neighbors, KNN), and CD4 and CD8A RNA expression are shown overlaid (right). Proportions of cells from each donor and assigned to each CAR are shown as pie charts. (C) Normalized expression of the top 50 differentially expressed genes within each KNN cluster represented in a heatmap. Significant differential expression was determined by a Wilcoxon rank-sum test with Bonferroni correction (adjusted *P* &lt; 0.05). If genes were present in multiple clusters, they are only displayed once. (D) Heatmap of chi-squared Pearson residual values representing enrichment of each CAR in the library within each KNN cluster. (E) Normalized expression of markers of T cell memory/stemness (left) and T cell activation and effector function (right). S.Score represents a metric of cell cycle activity, as implemented in the Seurat function, CellCycleScoring. Cluster 6, significantly enriched in cells expressing CAR1, is highlighted in red.

We collected more than 35,000 single-cell transcriptomes that could be confidently assigned to an individual CAR. To identify any potential donor-specific effects, the experiment was split between T cells derived from two healthy individuals. We achieved close to equal coverage of each CAR variant in each donor; following batch correction, dimensional reduction, and unsupervised clustering, these cells fell into nine different transcriptional clusters (**Figure 2B** and **Figure S2E-F**). Although CD4 and CD8 expression was differentially enriched in some clusters (**Figure S2G-H**), clustering is driven mostly by various biologically relevant markers of T cell activation (*IL2RA, HLA-DPA1, IFNG*), proliferation (*MKI67*, *TOP2A*), cytotoxicity (*GZMA, GZMB*), memory (*CCR7, SELL*), and stemness (*TCF7, IL7R, LEF1*) (**Figure 2C**). To assess how CAR ICD composition affects these cell states, we examined the distribution of each CAR variant within each cluster. As measured by chi-squared residuals, numerous CARs are enriched in specific clusters compared to what would be expected by random chance (**Figure 2D**), suggesting that different signaling inputs induce distinct transcriptional programs. As an initial validation of this hypothesis, we analyzed the gene signature associated with CAR1; lacking functional ICDs, CAR1-expressing cells should exhibit an unstimulated transcriptional phenotype. Consistent with this hypothesis, we found that CAR1 is strongly enriched only in cluster 6, which is characterized by significant overexpression of naïve T cell markers, as well as reduced expression of genes associated with activation, cytotoxicity, and proliferation (**Figure 2E**). Taken together, these results demonstrate that our ICD library induces a diversity of biologically relevant cell states, which we have captured in a robust transcriptional dataset mapping each signaling input to a defined molecular output.

### Defining classes of transcriptional programs activated by different CAR signals

Hypothesizing that diversity in input signals drives distinct transcriptional programs, we next sought to classify library variants based on similarities and differences in gene regulatory network (GRN) activity. To approach this in a data-driven manner, we identified transcription factor (TF)- target GRNs within our dataset based on modules of co-expressed genes and known sequence motifs of TFs, as implemented specifically for scRNA-seq data in the SCENIC algorithm.^33,34^ In brief, we utilize multiple runs of the SCENIC algorithm on our dataset to identify robust GRNs and score individual cells on the basis of GRN activity (**Table S4**; see **Methods**), generate psuedobulked scores that represent the average GRN activity for each CAR variant, and group each CAR into classes on the basis of these scores using k-means clustering (**Figure 3A**). The output of this analysis on aggregated data from both donors is shown in **Figure 3B**, allowing delineation of CAR variants into seven different classes, k1-k7, as measured by activity across 121 identified GRNs. As expected, CAR1 falls into its own class (k1), characterized by high activity of the LEF1 GRN, which is consistent with the downregulation of LEF1 following T cell activation.^35^ The remaining classes were labeled by increasing distance from CAR1 (**Figure S3A-C**), with k2 and k3 being most similar and k7 most different. Notably, each class exhibits differential activity of many TFs implicated in CAR T cell function, including the AP-1 family (BATF, JUNB),^36–38^ STATs (STAT2, STAT6),^17^ FOXO1,^39^ and RUNX3,^40^ suggesting that these CAR classes could induce transcriptional programs capable of driving differential phenotype and function.

**Figure 3.**
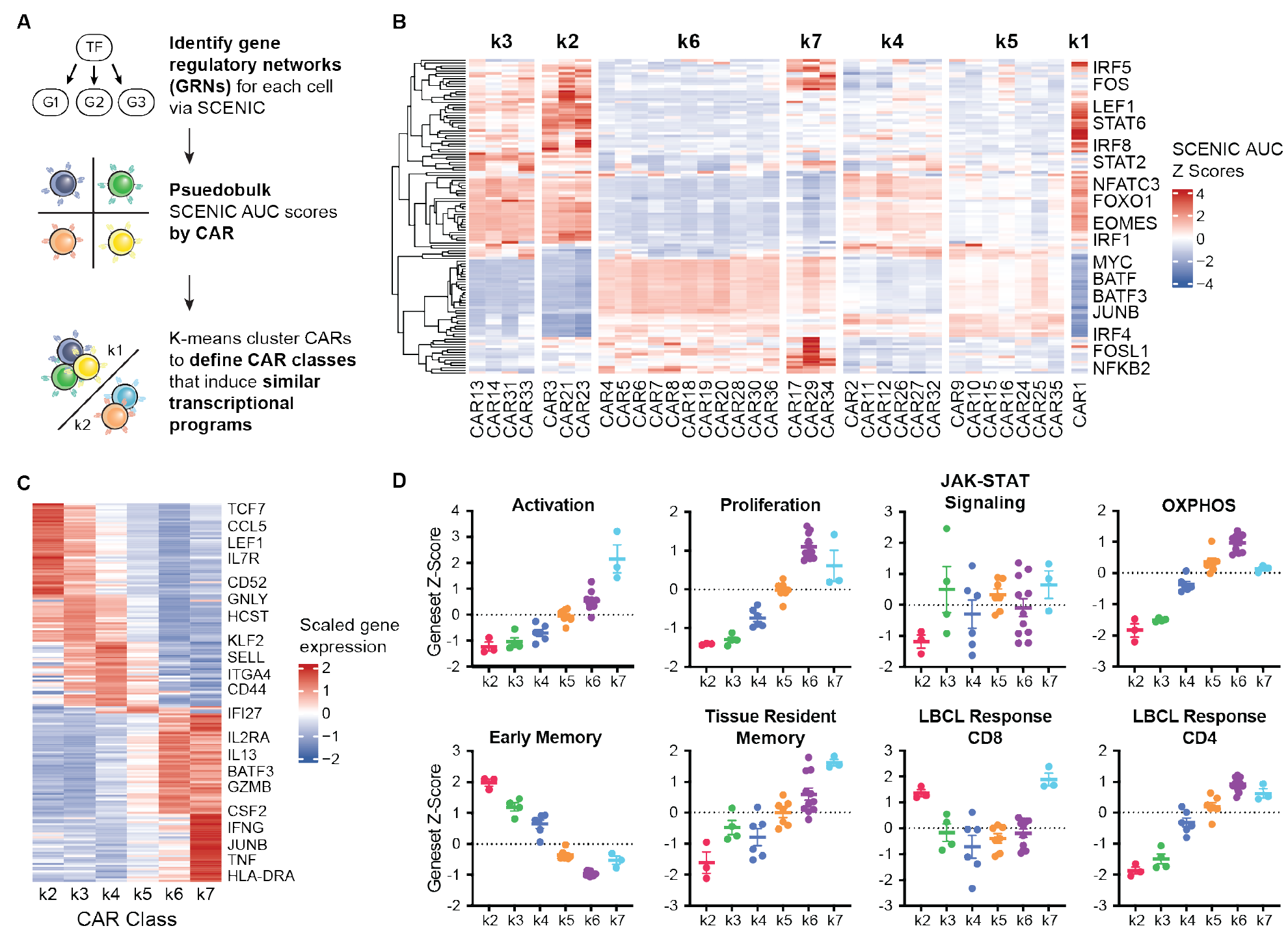
Defining classes of ICD architectures that induce distinct transcriptional programs. (A) Analysis pipeline to identify classes of CAR variants that induce similar transcriptional programs from the dataset in **Figure 2**. Gene regulatory networks (GRNs) are identified at the single-cell level using the SCENIC algorithm,^33^ before pseudobulking by CAR, and finally k- means clustering similar CARs by GRN activity. (B) Heatmap depicting SCENIC AUC scores pseudobulked by CAR, represented as *z-*scores, across all identified SCENIC GRNs. CARs are shown stratified into seven classes (k1-k7), identified by k-means clustering on pseudobulked SCENIC *z-*scores. Dendrograms represent hierarchical clustering of GRNs. (C) Normalized expression of the top 50 differentially expressed genes within each CAR class represented in a heatmap. Significant differential expression was determined by a Wilcoxon rank-sum test with Bonferroni correction (adjusted *P* &lt; 0.05). If less than 50 significant genes were detected for a given class, only significant genes are shown. If genes were detected across multiple classes, they are only displayed once. (D) Geneset *z-*scores for eight different immunologically relevant pathways, averaged across k- means classes. Scores were calculated via the *UCell* algorithm.^56^ Data points represent pseudobulked geneset scores for each CAR within each class. The mean ± SEM across each class is presented.

To further contextualize these classes, we also analyzed the signatures that distinguish them at both the gene and pathway level. Consistent with its similarity in GRN activity, k2—and to a lesser extent, k3—overexpresses similar gene markers as that of CAR1 (**Figure 3C** and **Figure 2E**), such as *TCF7, LEF1,* and *IL7R*. This is likely indicative of repeated antigen stimulation driving a state of dysfunction that resembles inactivation; all CARs that lost tumor control by day 7 fall into k2 or k3 (see **Figure S2A**). In contrast, the other classes show robust upregulation of different activation markers, such as *CD44*, *IL2RA*, and *HLA-DRA*, as well as effector molecules like *GZMB, IFNG,* and *TNF*. Relative expression of these genes, along with downregulation of memory markers like *SELL,* varies across classes, suggesting that these CAR classes might correspond to a spectrum of increasing T cell activation states and effector differentiation. These results are consistent with measurements at the pathway level, using a series of immunologically relevant literature-derived gene sets (**Table S5**), where we observe increasing activity of a CAR T cell activation module^10^ from k2 through k7 (**Figure 3D**). Other gene set scores reflect distinctions in the metabolism of each class, levels of JAK-STAT signaling, and T cell memory state. Notably, we also observe different levels of expression in gene signatures associated with patient response in large B cell lymphoma (LBCL),^41^ which lends clinical relevance to these delineations. In sum, these results—attained using separate gene sets from those identified by our GRN-based approach—indicate strong distinctions between classes and strengthen our CAR classifications.

Finally, examining the ICD architectures that comprise each class reveals some interesting patterns (**Figure S3D**). First, although most costimulatory domains appear to drive membership in particular classes to some extent—for example, 2B4 appears to exclusively direct cells to a k3- like state, regardless of the presence of DNAM1—the presence of other signals can provide confounding effects. One example of this is CD40—which appears to mainly exist in k7 (CAR29, MYD88.CD40.CD3z and CAR34, CD40.CD3e ITAM.DAP12), except in the context of CD79A, an architecture associated with k6 (CAR28, CD79A.CD40.CD3z). Another example is ICOS—which is either represented in k4 (CAR11, ICOS.CD3z) or k2 (CAR21, CD28.ICOS.CD3z and CAR23, 4-1BB.ICOS.CD3z), depending on the presence of an additional costimulatory domain. Second, we find that 28z (CAR4) and BBz (CAR5)—the two most common second-generation CARs used in the clinic that are known to drive distinct phenotypes—group together in k6. Although our dataset does recapitulate the significant upregulation of genes known to distinguish BBz from 28z when they are directly compared (**Figure S3E**),^10^ this difference is small relative to the rest of the library; indeed, this same gene signature is upregulated to a much greater degree in k7 constructs bearing CD40. These results suggest that the CARs currently in clinical use are greatly undersampling the diversity of cell states available to synthetically programmed T cells.

### Functional characterization of CAR candidates across CAR classes

Given the transcriptional distinctions observed between CAR classes, we selected a single candidate CAR from each class to thoroughly link transcriptional states, molecular responses, and phenotypic outcomes. To try to maximize phenotypic diversity, we identified a pool of candidates with the highest Euclidean distance from each other based on GRN activity, while avoiding those that failed to control NALM6 cells over the one-week rechallenge. From k6, we selected CAR5 (BBz) to ensure we included a clinical benchmark. Seven candidates were chosen, covering a range of different ICDs in various contexts: CAR1 (k1, non-signaling control), CAR21 (k2, CD28.ICOS.CD3z), CAR33 (k3, DNAM1.2B4.CD3z), CAR12 (k4, DAP10.CD3z), CAR25 (k5, CD28.CD3z.DAP10), CAR5 (k6, 4-1BB.CD3z), and CAR29 (k7, MYD88.CD40.CD3z). Each candidate was challenged with NALM6 cells *in vitro* before assessing cytotoxic capacity by live cell imaging, cytokine secretion by ELISA, and memory state by flow cytometry. While the candidates appear largely indistinguishable when challenged with a low tumor burden (effector:target ratio, E:T = 1:1) (**Figure S4A**), under high tumor burden challenges (E:T = 1:10), we found that those from k2 and k3 lose control over NALM6 growth over time, those from k4-k6 exhibit intermediate killing, and the k7 candidate allows the least tumor growth (**Figure 4A**). Similarly, we observed increasing levels of IL-2, IFNγ, and TNFα secretion from k2-k7 (**Figure 4B**), which may mediate enhanced killing. These observations are consistent with the increasing levels of activation we saw in transcriptional signatures across classes. Interestingly, we found that this increased effector function does not drive greater amounts of terminal differentiation following a week of antigen stimulation (**Figure S4B**), with all candidates showing similar levels of CD62L^−^CD45RA^+^ (TEMRA) subsets.

**Figure 4.**
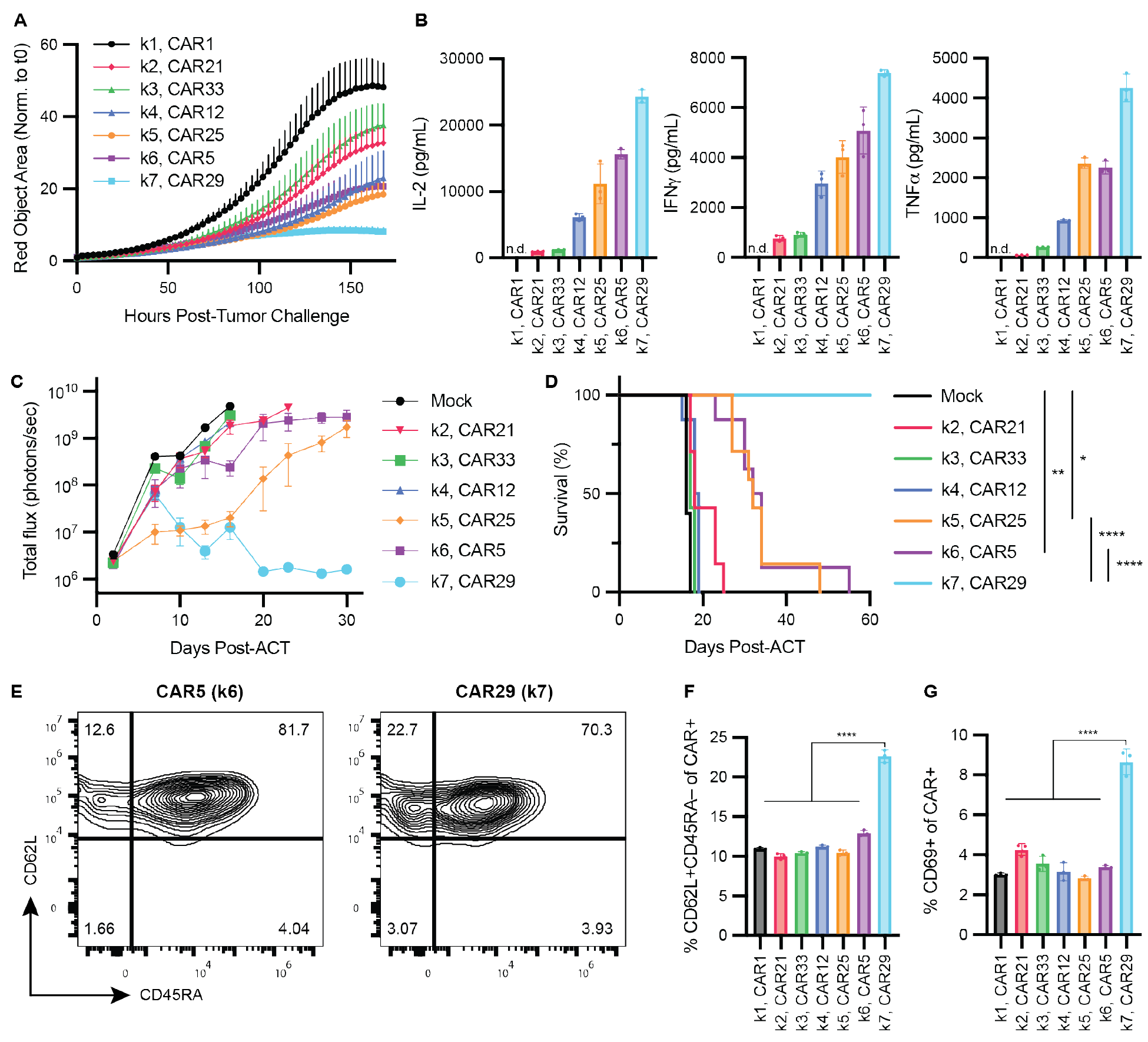
Candidates from each class exhibit significant functional differences both *in vitro* and *in vivo*, under antigen stimulation, and at rest. (A-B) Functional characterization of candidate CARs from each CAR class identified in Figure 3, following NALM6 tumor challenge (E:T = 1:10). Longitudinal control of tumor burden over seven days, as measured by live cell imaging, is shown in (A), and cytokine secretion levels at 48h post-stimulation, as measured by ELISA, are shown in (B). (C-D) Characterization of *in vivo* efficacy driven by candidate CARs (*n* = 7-8 mice per group). Mice bearing NALM6 tumors were treated at a dose sub-curative for a 19BBz/CAR5 control CAR (2x10^5^ CAR^+^ cells), and tumor burden was tracked longitudinally by bioluminescence. Tumor growth curves are shown in (C), while overall survival is shown in (D). *P* values were calculated by log-rank Mantel-Cox test, prior to Bonferroni multiple hypothesis correction using all possible pairwise comparisons (\**P* &lt; 0.05; \*\**P* &lt; 0.01; \*\*\*\**P* &lt; 0.0001). (E-G) Functional characterization of candidate CARs at rest. Representative flow plots showing expression of memory surface markers for k6 and k7 candidates in the absence of antigen are shown in (E) and quantified across all CARs in (F) as the percentage of CD62L^+^CD45RA^−^ CARs. Expression of the activation marker CD69 at rest is quantified in (G). *P* values were determined by one-way ANOVA with Tukey’s multiple comparisons test (\*\*\*\**P &lt;* 0.0001). All pooled data is shown as mean ± SEM. *In vitro* studies in (A-B, E-G) show three technical replicates from a representative donor.

Motivated by these functional distinctions observed *in vitro*, we characterized each candidate *in vivo* to elucidate the impacts of these different phenotypes on anti-tumor efficacy. To this end, we treated NALM6 tumor-bearing mice with each candidate CAR using a dose that is subcurative for BBz/CAR5 (2x10^5^ CAR^+^ cells), allowing us to distinguish any candidate CARs with enhanced efficacy compared to the clinical state-of-the-art. The different CARs strongly stratified mice into non-responders (NR), in which k2-k4 candidates were indistinguishable from mock-transduced T cells; partial responders (PR), in which k5-k6 candidates significantly delayed tumor growth but eventually lost tumor control; and complete responders (CR), in which mice treated with the k7 candidate (CAR29) remained tumor free for the entirety of the two-month study (**Figure 4C-D**). The potent anti-tumor response induced by CAR29 was seemingly driven by long-term persistence of CAR T cells, as spleens from CAR29-treated mice retain significantly higher amounts of T cells *in vivo* compared to PR and NR groups as early as day 16 post-ACT (**Figure S4C**). Strikingly, similar numbers of CAR29 cells could be detected as long as 56 days after treatment, following more than a month of remission (**Figure S4D**).

We hypothesized that the distinctions in *in vivo* persistence that we observed across CAR candidates might be due to differences in tonic signaling, as baseline levels of activation are thought to impact exhaustion and persistence.^36,42–44^ To address this, we examined cell states of each candidate in the absence of antigen stimulation. This analysis revealed striking differences between the k7 candidate and the others, indicating increased levels of tonic signaling. Looking first at memory surface markers at rest, we found that cells expressing k1-k6 candidates are indistinguishable, but CAR29 enriched for a central memory phenotype, driving a significantly higher proportion of CD62L^+^CD45RA^−^ cells (**Figure 4E-F**). We observed similar patterns in activation marker expression at rest, with CAR29 inducing CD69 expression at a much higher rate than the other candidates (**Figure 4G**). Taken together, functional characterizations of candidate CARs indicate that the transcriptional programs induced by k7 CARs induce strong acute activation and effector function following antigen challenge, elicit tonic signaling at rest, and enhance *in vivo* persistence, all of which aid in complete long-term cures in this liquid tumor model. Notably, these *in vitro* and *in vivo* results were recapitulated using CAR T cells derived from a different donor (**Figure S4E-K**), suggesting variability between individuals is not significantly affecting differences in CAR function.

### Defining gene expression profiles associated with liquid tumor efficacy

Given the strong *in vivo* efficacy profile exhibited by the k7 CAR candidate, we returned to the gene expression dataset to ask whether we could identify molecular correlates of CR in this model. We first utilized partial least squares regression (PLSR)^45^ to detect how GRN activity scores affected *in vivo* survival (see **Methods**). PLSR yielded one component (41% variance) that separated k7 CARs from others (**Figure 5A**), and the activity of a subset of GRNs demonstrated varying associations with survival (**Figure 5B**). Of those positively associated with efficacy, many TFs have been implicated in T cell activation, memory, and persistence (*NFKB2*,^46^ *RXRA*,^47^ and *AHR*^48^). We also identify members of the AP-1 TF family (*FOSL2, JUN*) associated with efficacy, both of which have been shown to enhance *in vivo* persistence and efficacy when overexpressed in engineered T cells.^36,49^ As an orthogonal approach, we also used differential gene expression analysis comparing k7 CARs to those that exhibit a PR (k5 and k6). This revealed numerous significantly upregulated genes, including inflammatory cytokines such as *IFNG, TNF, CSF2, CXCL10, CCL3, CCL4,* and *XCL1*; various elements of the NF-kB pathway (*NFKB2, NFKBIA,* and *TRAF1*) and *JUNB*; and various genes related to class II MHC expression, such as *CD74, HLA-DRA*, and *HLA-DPA1* (**Figure 5C** and **Table S6**). These observations are consistent with the PLSR results, which identified *RFX2*—known to regulate class II MHC expression^50^—*NFKB2*, and AP-1 TFs as positively associated with efficacy. Cytokine overexpression also aligns with increased TNFα and IFNγ secretion measured during candidate phenotyping. To contextualize these genes biologically, we performed gene set enrichment analysis (GSEA) against a plethora of canonical pathways, finding significant enrichment of NF-kB signaling and various inflammatory cytokine-related pathways (**Figure 5D**). Interestingly, we also observe enrichment of the CD40- CD40L signaling pathway, which might be indicative of a dominant role of the CD40 ICD in 2/3 k7 CAR architectures. These results also point to a metabolic switch between CR and PR CARs, with glycolysis enriched in k7 and oxidative phosphorylation enriched in k5/k6—this is a particularly interesting result given previous reports that glycolytic metabolism in CD28-based CARs elicits decreased persistence,^9^ which we do not observe in k7 CARs.

**Figure 5.**
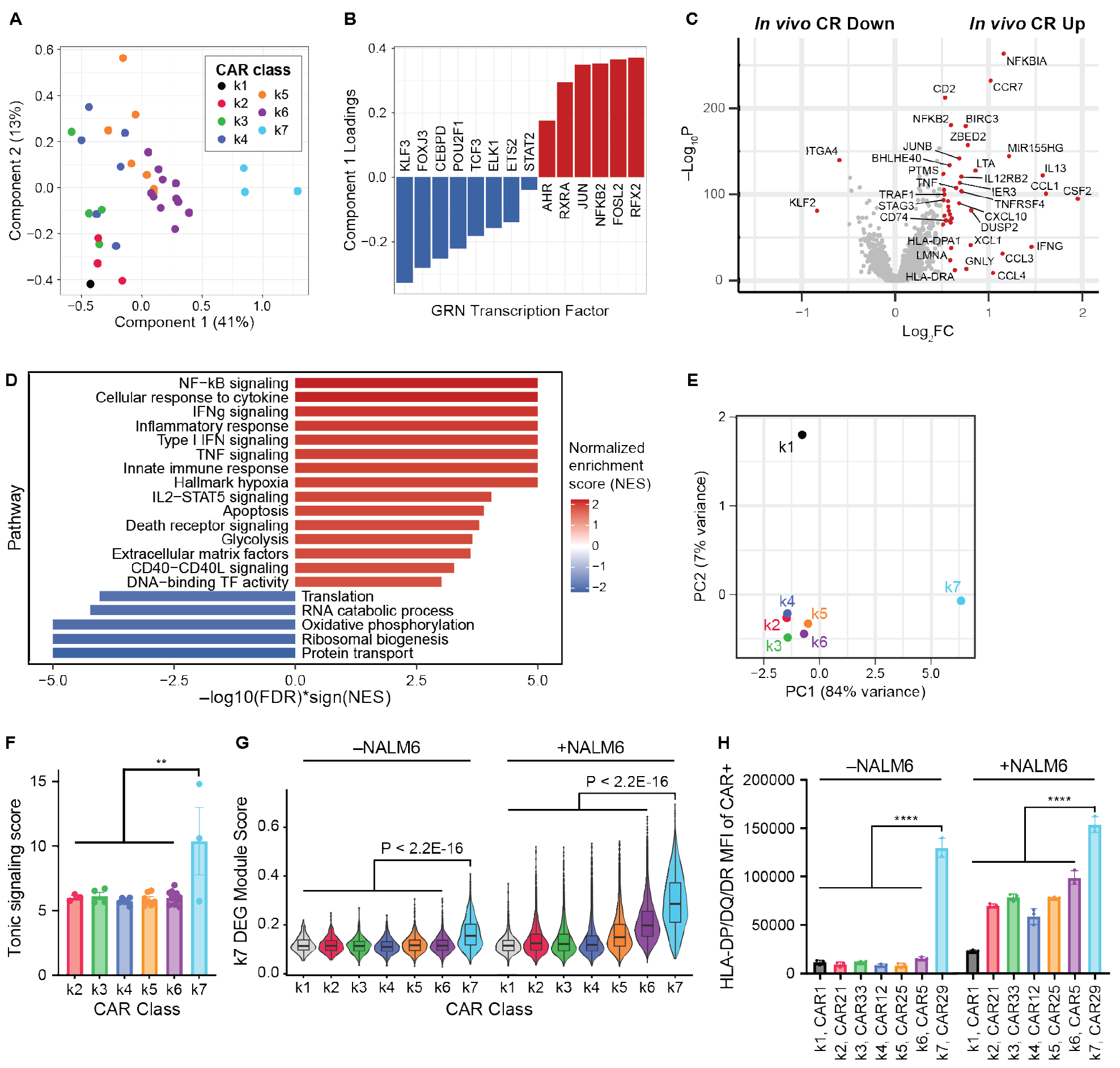
Definition of an ICD-intrinsic gene expression profile associated with durable *in vivo* efficacy. (A-B) Partial least squares regression (PLSR) results in which GRN activity scores were regressed on NALM6 *in vivo* survival time. Transformed GRN scores for each CAR are displayed in the resulting latent space in (A), plotted across the first two components. Each CAR is colored by CAR class. Component 1 loadings for GRNs in the PLSR model are shown in (B). (C) Volcano plot showing differentially expressed genes between k7 (complete responders, CR) to k5 and k6 (partial responders) cells from the dataset introduced in **Figure 2**. Unadjusted *P* values from a Wilcoxon rank-sum test are plotted. Genes are highlighted in red with Log2FC > 0.5, and with Bonferroni-adjusted *P* &lt; 0.05. (D) Pathway enrichment analysis via gene set enrichment analysis (GSEA) of the differentially expressed genes shown in (C). Pathways selected from those with FDR &lt; 0.001 are shown. (E) Transcriptional characterization of CAR classes measured at rest, in the absence of antigen, depicted across the first two principal components (PCs), which were calculated using all variable genes, pseudobulked across all cells expressing CARs within each class. (F) Tonic signaling scores calculated across CAR classes, computed as the Euclidean distance from each CAR to CAR1 in the absence of antigen, across all variable genes. Each datapoint represents a single pseudobulked CAR within each class. (G) Module scores representing expression of the k7 signature in the absence (resting dataset) and presence (antigen rechallenged dataset) of antigen, as defined by the top 50 differentially expressed genes (DEGs) shown in (C), across all cells, grouped by CAR class. Scores were calculated by the *UCell* algorithm.^56^ *P* values were calculated between each group within each dataset by Wilcoxon rank-sum test. (H) Protein-level validation of a k7 marker gene, HLA class II, measured both in the absence and presence of antigen (NALM6 cells at E:T = 1:1, n = 3 technical replicates). Expression was quantified as mean fluorescence intensity (MFI), measured via flow cytometry. *P* values were determined by one-way ANOVA with Tukey’s multiple comparisons test (*\*\*\*\*P* &lt; 0.0001). Pooled data in F and H are shown as mean ± SEM.

The k7 candidate (CAR29) also exhibited increased tonic signaling at rest, which may have an impact on efficacy. To confirm this effect and investigate its molecular basis, we performed caRNA-seq on the library in the absence of antigen, again achieving approximately equal coverage of all CAR variants in the library (**Figure S5A**). Direct comparison of the rested library against the antigen rechallenged library demonstrates significantly reduced expression of effector and activation molecules, along with overexpression of naïve T cell markers (**Figure S5B**), as expected. Looking at pseudobulked gene expression profiles across CAR classes in this dataset, we observed a stark distinction between k7 CARs and the other classes, with k7 sitting alone in principal component (PC) space (**Figure 5E**). To assess levels of tonic signaling across the library, we calculated a tonic signaling score based on the distance between each CAR and CAR1 in gene expression space—given its lack of functioning ICDs, we reasoned that any deviation from CAR1 would be due to tonic signaling. By this analysis, all other classes scored similarly, while the k7 class scored significantly higher (**Figure 5F**). Notably, this effect was dominated by the two CD40-based CARs in k7, potentially suggesting a CD40-mediated impact on tonic signaling (**Figure S5C**). To validate this finding against an orthogonal measure of tonic signaling, we also scored each CAR on the basis of a recently published gene set associated with CD3z-mediated activation in the absence of antigen;^10^ consistent with our scoring, the k7 class exhibits higher expression of this tonic signaling module (**Figure S5D**). Interestingly, the same set of marker genes that distinguish k7 CARs from k5/k6 CARs in the antigen rechallenged dataset remain upregulated exclusively in k7 CARs in this dataset (**Figure 5G**), suggesting that the same transcriptional programs that distinguish k7 after antigen challenge are also activated tonically. To confirm these observations at the protein level, we assessed MHC class II expression—inclusive of several marker genes upregulated in k7— across CAR candidates, in the presence and absence of antigen. These data confirmed significant upregulation of HLA-DP/DQ/DR exclusively in CAR29, both before and after antigen challenge (**Figure 5H**). Collectively, these data, all of which are consistent with our previous candidate phenotyping, provide strong evidence—at both the transcriptional and protein level—that increased levels of tonic signaling distinguish k7 CARs from the other CARs in the library.

### Characterizing functional and transcriptional shifts in response to solid tumor perturbation

Having characterized our library in a liquid tumor model, we next set out to translate it to a solid tumor model given the poor efficacy of CAR T cells in solid tumors. To do this without changing the CAR backbone, we utilized a melanoma cell line, A375, with ectopic expression of CD19. Exposing our CAR candidates to *in vitro* A375 challenge, we recapitulated the results seen in NALM6, with increasing cytotoxic capacity from k2-k7 candidate CARs (**Figure 6A**), as well as increasing inflammatory cytokine secretion (**Figure 6B**). However, in a comparable subcutaneous xenograft model using CD19^+^ A375, again dosing with a subcurative dose for CAR5 (5x10^5^ CAR^+^ cells), the *in vivo* efficacy changed dramatically compared to what we observed in NALM6. In this context, only k5 and k6 candidates demonstrated any statistically significant survival benefit compared to mice treated with mock-transduced T cells (**Figure 6C-D**). In stark contrast to NALM6, CAR29-treated mice failed to control *in vivo* tumor growth.

**Figure 6.**
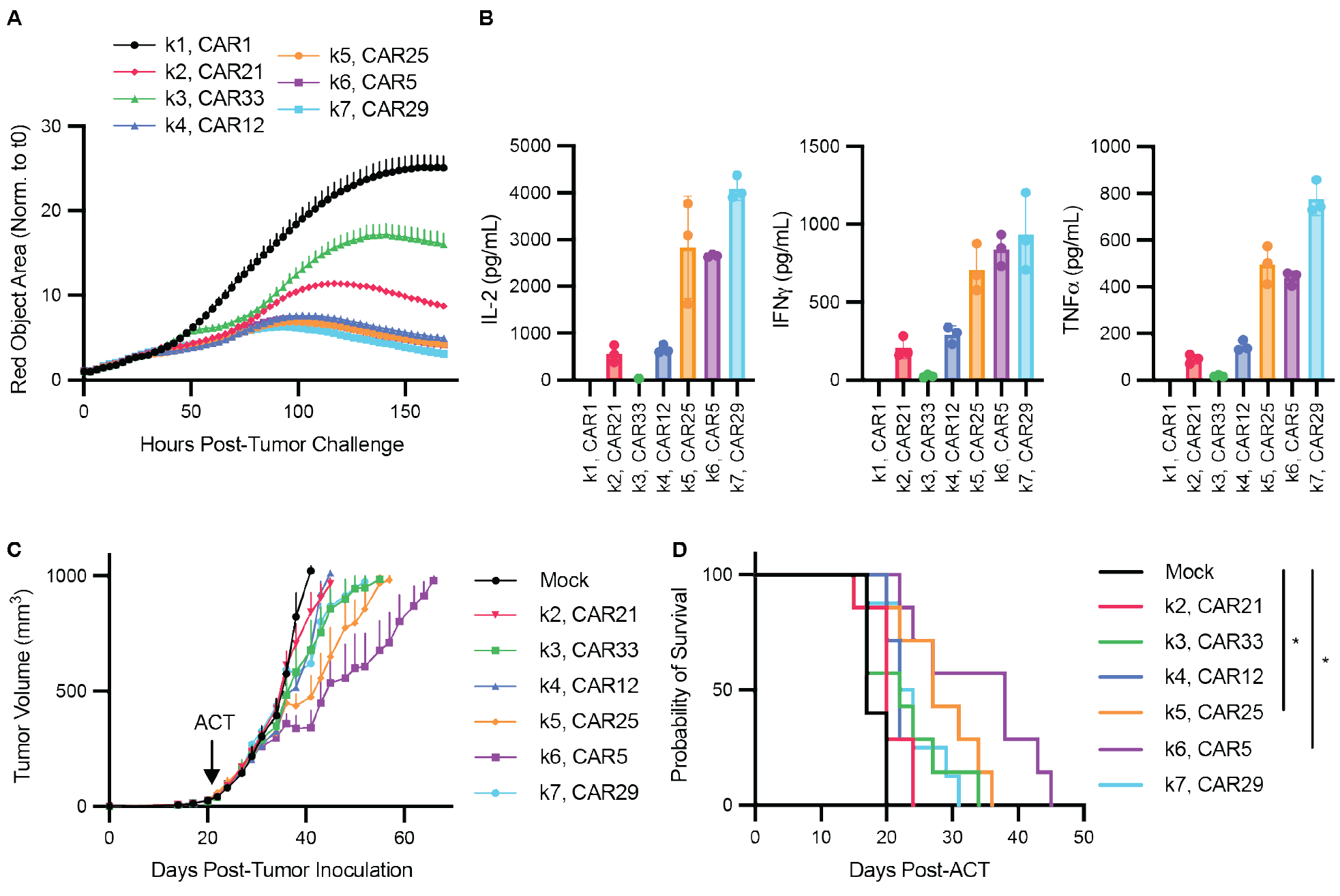
CAR candidates challenged with solid tumors exhibit similar *in vitro* function as in liquid tumors but differ greatly *in vivo*. (A-B) Functional characterization of candidate CARs from each CAR class identified in Figure 3, following CD19^+^ A375 tumor challenge (E:T = 1:10). Longitudinal control of tumor burden over seven days, as measured by live cell imaging, is shown in (A), and cytokine secretion levels at 48h post-stimulation, as measured by ELISA, are shown in (B). (C-D) Characterization of *in vivo* efficacy driven by candidate CARs (n = 7-8 mice per group). Mice bearing A375 tumors were treated at a dose sub-curative for a 19BBz/CAR5 control CAR (5x10^5^ CAR^+^ cells), and tumor size was measured longitudinally. Tumor growth curves are shown in (C), while overall survival is shown in (D). *P* values were calculated by log-rank Mantel-Cox test, prior to Bonferroni multiple hypothesis correction using all possible pairwise comparisons (\**P* &lt; 0.05). All pooled data is shown as mean ± SEM. *In vitro* studies in (A-B) show three technical replicates from a representative donor.

Based on the inconsistencies we observed between *in vitro* and *in vivo* A375 models, we hypothesized that the *in vivo* solid tumor microenvironment might elicit a different signaling landscape than what we observed *in vitro.* To address this question, we performed caRNA-seq directly on library-treated *in vivo* solid tumor samples. We treated A375-bearing mice with 1x10^6^ cells expressing the CAR library in pool before sorting and sequencing CAR T cells harvested from the tumor of three different mice, which were separately hashed with unique CITE-seq antibodies to allow deconvolution after sequencing (**Figure 7A**). In addition to hashing antibodies, we also stained with a panel of CITE-seq antibodies against several T cell markers. Tissue was harvested 10 days post-treatment, after the CAR-treated tumors had deviated from mock-treated controls, suggesting a CAR-mediated response was ongoing (**Figure 7B**). Initial characterization of the library as a whole revealed six transcriptional states split between CD4 and CD8 T cells, which varied in expression of cytotoxic genes, cytokines, and activation markers, as well as those indicative of T cell memory and exhaustion (**Figure 7C** and **Figure S6A-B**). Collectively these findings demonstrate that we have captured an active anti-tumor response. Importantly, cells harvested from different mice appear indistinguishable in UMAP space, indicating strong consistency in transcriptional response between replicates and allowing us to aggregate data across mice.

**Figure 7.**
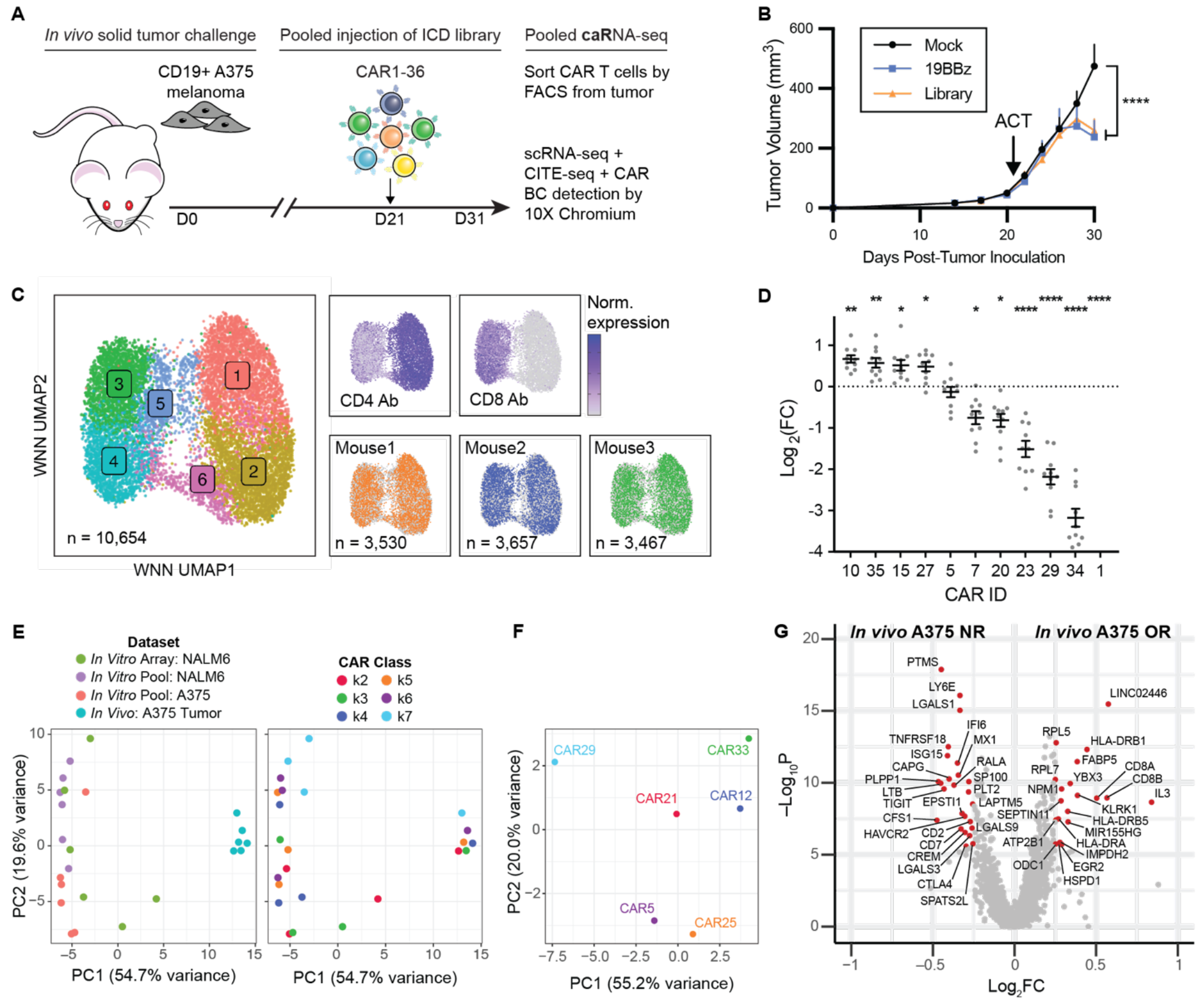
Transcriptional characterization of the library under *in vivo* solid tumor challenge via caRNA-seq reveals a marked shift in signaling response. (A) Experimental design to assess transcriptional profiles under *in vivo* solid tumor challenge via caRNA-seq. Tumor-bearing mice were treated with 1x10^6^ library-expressing CARs in pool. Ten days post-treatment, CAR T cells were sorted from the tumor for caRNA-seq. Tumors were stained with CITE-seq antibodies specific for T cell phenotypic markers. (B) Tumor growth curves prior to CAR T cell harvest. *P* values were calculated by two-way ANOVA with Dunnett’s multiple comparisons correction (\*\*\*\**P* &lt; 0.0001). (C) UMAP embeddings of all cells harvested from the tumor that were confidently called to a single CAR variant (*n =* 10,654 cells). Weighted-nearest neighbor analysis (WNN) was used to incorporate CITE-seq antibody data into the dimensional reduction and unsupervised clustering results depicted on the left. CD4 and CD8 CITE-seq antibody expression, as well as the mouse of origin, are shown overlaid (right). (D) Differential CAR enrichment within the tumor, as measured by fold change from pre-injection proportions, as measured by next-generation sequencing (NGS) of CAR BCs, to those observed at tumor harvest. Data points represent different tumors from individual mice, compiling both NGS (*n =* 7 mice) and scRNA-seq cell proportions (*n =* 3 mice). *P* values were determined by one-way ANOVA with Tukey’s multiple comparisons test (\*\*\*\**P &lt;* 0.0001), where each CAR was compared to the enrichment observed in the CAR5/19BBz control. Only CARs with significantly different enrichment levels are shown. (E) Principal component analysis (PCA) results from scoring different CAR classes across contexts and tumor models on gene regulatory network (GRN) activity, using the GRNs identified in **Figure 3**. Data points represent pseudobulked GRN activity for cells expressing CARs in each class in four different contexts: arrayed *in vitro* NALM6 rechallenge, pooled *in vitro* NALM6 rechallenge, pooled *in vitro* A375 rechallenge, and *in vivo* A375 solid tumor. Groups are colored by dataset (left) or CAR class (right). (F) PCA of a subset of *in vivo* solid tumor dataset, only considering candidate CARs with matched phenotyping data in **Figure 6**. Datapoints represent pseudobulked expression of variable genes averaged across cells expressing each CAR. (G) Volcano plot showing differentially expressed genes between responding CARs (CAR5 and 25) and non-responding CARs (CAR12, 21, 29, and 33). Unadjusted *P* values from a Wilcoxon rank-sum test are plotted. Genes are highlighted in red with Log2FC > 0.25, and with Bonferroni-adjusted *P* &lt; 0.05. Pooled data in (B) and (D) is presented as mean ± SEM.

To explore ICD-specific responses within this dataset, we first analyzed relative enrichment of each CAR variant within the tumor across 10 different mice, utilizing a combination of cell proportions detected in the scRNA-seq dataset (n = 3 mice), short-read sequencing of CAR BCs from tumor-derived genomic DNA (n = 7 mice), as well as amplicon sequencing of the pre-injection pool to define the initial library composition. As expected, CAR1 was the least frequent variant detected in the tumor; interestingly, CAR29 and CAR34, the two CD40-based k7 CARs, were also significantly de-enriched relative to the CAR5 benchmark (**Figure 7D** and **Figure S6C**). This suggests that CAR29-expressing cells poorly infiltrate, expand, and/or persist within the *in vivo* tumor microenvironment, which likely contributes to the poor efficacy we observed in candidate phenotyping.

To assess how ICD-intrinsic transcriptional programs shift from the *in vitro* setting to that of the tumor microenvironment, we next calculated GRN activity across tumor-infiltrating CARs, using the same GRNs identified *in vitro*. We also generated matched datasets under pooled *in vitro* rechallenges in both NALM6 and A375 models to isolate effects to the *in vivo* environment, rather than the effect of CAR-CAR crosstalk or the A375 model. Comparing GRN scores across CAR classes in each setting via PCA, we found that the *in vivo* context was the major driver of variability in GRN activity, with PC1 (54.7% of variance) strongly separating all *in vitro* samples from *in vivo* (**Figure 7E**). Notably, all k7 *in vitro* samples cluster together, regardless of context, but this does not translate *in vivo*. These results indicate a distinct shift in CAR-intrinsic transcriptional programming from what we observe *in vitro*, likely mediated by the solid tumor microenvironment. In addition, the CAR classes assigned *in vitro* are largely indistinguishable in the *in vivo* solid tumor, suggesting that our library variants may not fall into the same groupings following these transcriptional shifts.

Given these discrepancies between *in vitro* and *in vivo* transcriptional programs, we focused our remaining analysis specifically on those candidate CARs for which we had matched efficacy phenotyping data, rather than binning by GRN classes identified *in vitro.* Pseudobulk profiles of only these CARs separate cleanly into responders (CAR5 and 25) and non-responders (CAR12, 21, 29, and 33) (**Figure 7F**). Analysis of the markers that distinguish these two groups revealed overexpression of several exhaustion markers (*HAVCR2, TIGIT, CTLA4*) in non-responding CARs (**Figure 7G** and **Table S7**), suggesting that these particular ICD architectures are prone to exhaustion when faced with the solid tumor microenvironment. On the other hand, responding CARs upregulate activation markers, such as *MIR155HG* and a suite of MHC class II molecules (e.g. *HLA-DRA, HLA-DRB5, HLA-DRB1*), indicating that these CARs are likely directing the anti-tumor response observed at the time of harvest. Overexpression of class II MHC is notable, as this was associated solely with CAR29 in our other candidate phenotyping—this finding is again consistent with the hypothesis that ICD-intrinsic signaling responses are dramatically different in this solid tumor setting. Lastly, CD8 expression appears greatly enriched in responding CARs, suggesting that a CD8 skew may be beneficial in this context. These genes provide a potential signature associated with *in vivo* response against solid tumors, which could be useful in future ICD screening efforts.

## Discussion

Although there have been extensive engineering efforts to enhance CAR signaling, a systematic understanding of the structure-function relationships that dictate differential phenotype and function has remained elusive. To address this, we used caRNA-seq to systematically characterize diverse ICD architectures, yielding robust transcriptional datasets that separated CARs into seven distinct groups on the basis of GRN activity. Looking broadly at how ICDs are distributed across these classes, we observe some patterns indicative of specific signals driving particular transcriptional programs (e.g. 2B4); however, the majority of ICDs exist across multiple classifications (e.g. CD40, ICOS, CD27), which precludes identifying simple relationships between CAR ICD identity and downstream T cell phenotype. Instead, it has proven necessary to generate datasets capable of empirically decoding these relationships.

In addition, although we recapitulated known differences between 28z (CAR4) and BBz (CAR5) when compared directly, we found that they cluster together within k6 in the context of the library as a whole. This suggests that a plethora of other phenotypes are available within synthetically engineered T cells beyond those commonly used in the clinic. This is clear even within the limited scope of this defined library; it is likely that an even greater diversity of phenotypes—with potentially enhanced efficacy profiles—might be accessible within more diverse libraries of ICDs.

One of the CAR classes that we identified, k7, induced a unique transcriptional signature associated with strong T cell activation, inflammatory cytokine secretion, and durable *in vivo* persistence, driving long-term cures in a challenging liquid tumor mouse model. Interestingly, this class exhibited significantly greater levels of tonic signaling relative to the rest of the library. These findings are contrary to typical characterizations of tonic signaling, which, in excess, is thought to drive exhaustion and limit persistence;^36,42,43^ however, these observations are all in the context of CARs with canonical BBz and/or 28z ICDs. Under alternate signaling programs, it is possible that low levels of baseline activation could aid T cell survival and persistence, which would be consistent with the behavior of the k7 candidate that we chose for phenotyping. Notably, our results are largely consistent with the initial report of this construct (CAR29), which also showed strong levels of *in vivo* persistence along with baseline levels of activation in an EphA2-targeted CAR,^19^ indicating that these effects are applicable to different antigen binders. Others have also linked increased tonic signaling to enhanced efficacy,^44^ lending credence to our findings. Given the demonstrated correlation between persistence and long-term remission observed in patients,^51,52^ ICD architectures that maximize persistence—perhaps via this tonic signaling mechanism—could be strong candidates for clinical testing.

One of the main challenges facing CAR T therapies is their translation from liquid tumors to solid tumors, with poor responses observed in most solid tumor trials.^6^ In this study, we characterized our library under *in vivo* solid tumor challenge, using the same antigen target to allow direct comparisons between liquid and solid tumor contexts. To our knowledge, this is the first scRNA-seq dataset of its kind, in which a library of ICD architectures is characterized directly from a solid tumor. We observed marked differences in transcriptional responses within the solid tumor microenvironment compared to those *in vitro*, even within the same tumor model. This discordance was also observed functionally—CAR29 excelled under A375 *in vitro* challenge, but failed to control *in vivo* growth, in stark contrast to its durable control of NALM6. These findings highlight the significance of context in evaluating CAR architectures—the solid tumor microenvironment presents a unique set of molecular cues and physical barriers, which appears to induce distinct signaling responses in our datasets. Therefore, it stands to reason that the optimal ICD architecture for a liquid tumor may not be well suited for a solid tumor, as we observed with CAR29. In the clinic, however, the same sets of canonical ICDs are typically tested in solid and liquid tumors, highlighting the need for ICD engineering specifically for solid tumors.

Although we limited our investigations to only a single CD19-targeted backbone to eliminate any non-ICD-mediated effects, evaluating the transcriptional responses of ICD architectures across different target antigens, binding affinities, and CAR frameworks would further reveal CAR design principles. As one example, it is well demonstrated that CARs with different antigen-binding moieties can exhibit distinct tonic signaling profiles via mechanisms such as scFv dimerization;^7^ the anti-CD19 scFv (FMC63) used in our study induces relatively low levels compared to other binders.^36,53^ Similarly, analyses of other cell types under consideration as CAR therapies, such as NK cells and macrophages, may demonstrate altered ICD-intrinsic responses due to differences inherent to other immune cell lineages.

The therapeutic impact of CAR-T therapies continues to increase in cancer, and has recently extended to other disease indications such as autoimmunity and fibrosis.^54^ These diverse use cases may benefit from distinct T cell phenotypes, for which ICD architectures can be selected for each setting. To enable such targeted engineering will require robust datasets that span a diverse set of ICDs and disease contexts yet are amenable to robust comparisons, such as those presented here. In combination with recent advances in machine learning and computational modeling, which are beginning to be applied to CAR design^27^ and genome-wide transcriptional responses,^55^ predictive tools for the rational design of these next-generation CAR constructs are within reach.

## Acknowledgments

We thank the Koch Institute’s Robert A. Swanson (1969) Biotechnology Center for their technical support, especially the Flow Cytometry Facility, Preclinical Modeling, Imaging and Testing Core, and MIT BioMicro Center. M.E.B. was supported by a Packard Fellowship, a Pew-Stewart Scholarship and a grant from the Deshpande Center. This work was supported in part by the Koch Institute Frontier Research Program (to M.E.B.); the Bridge Project, a partnership between the Koch Institute for Integrative Cancer Research at MIT and the Dana-Farber/Harvard Cancer Center (M.E.B. and M.V.M); and the Koch Institute Support (core) Grant P30-CA14051 from the National Cancer Institute. We also acknowledge funding from the National Institute of Allergy and Infectious Diseases (contract #75N93019C00071). This work was additionally supported by a National Science Foundation Graduate Research Fellowship to C.R.P. and K.S.G., as well as a fellowship from the Ludwig Center at MIT’s Koch Institute to C.R.P. A.G. and B.S. were supported by award number T32GM144273 and T32GM007753, respectively, from the National Institute of General Medical Sciences. A.N. was supported by funding from the Swedish Research Council (2019-06349). The content is solely the responsibility of the authors and does not necessarily represent the official views of the National Institute of General Medical Sciences or the National Institutes of Health. This research is supported in part by the National Research Foundation, Prime Minister’s Office, Singapore under its Campus for Research Excellence and Technological Enterprise (CREATE) programme, through Singapore MIT Alliance for Research and Technology (SMART): Critical Analytics for Manufacturing Personalised-Medicine (CAMP) Inter-Disciplinary Research Group.

## Author Contributions

C.R.P and M.E.B. conceived the project. C.R.P., A.G., K.S.G., L.G.L., and B.E.S. conducted experiments. C.R.P., A.N., and H.M.B. performed data analysis. M.V.M, D.A.L., and M.E.B. supervised the work. C.R.P. and M.E.B wrote the manuscript. All authors contributed to the editing of the manuscript.

## Declaration of interests

M.E.B. is a founder, consultant, and equity holder of Kelonia Therapeutics and Abata Therapeutics. M.V.M. is an inventor on patents related to adoptive cell therapies, held by Massachusetts General Hospital (some licensed to Promab and Luminary) and University of Pennsylvania (some licensed to Novartis). M.V.M. receives Grant/Research support from Kite Pharma and Moderna, serves as a consultant for multiple companies involved in cell therapies, holds equity in 2SeventyBio, A2Bio, Affyimmune, BendBio, Cargo, Century Therapeutics, Neximmune, Oncternal, and TCR2 and serves on the Board of Directors of 2Seventy Bio. K.S.G. is currently employed at Ginkgo Bioworks, Inc. The other authors declare no competing interests.

## Methods

### Cell lines

HEK293T, Jurkat (clone E6-1), NALM6, and A375 cell lines were purchased from ATCC, and cultured in DMEM (ATCC) + 10% FBS + 100 U/mL penicillin/streptomycin (P/S) (HEK293T and A375) or RPMI-1640 (ATCC) + 10% FBS + 100 U/mL P/S (Jurkat and NALM6). Cells were maintained in normal cell culture conditions, at 37°C in 5% CO2. NALM6 cells for *in vivo* studies were transduced to stably express a firefly luciferase (FFLuc) gene via lentiviral transduction. To generate CD19^+^ A375 cells, a truncated CD19 ectodomain was introduced via lentiviral transduction, and transduced cells were sorted for comparable levels of CD19 expression as NALM6 cells, as described previously.^26^ For live cell imaging studies, mCherry expression was introduced to NALM6 cells and CD19^+^ A375 cells via lentiviral transduction. All cell lines were routinely tested for mycoplasma contamination (MycoAlert PLUS Mycoplasma Detection Kit, Lonza).

### Mouse strains

All animal studies in this study were completed under an approved International Animal Care and Use Committee protocol, following guidelines provided by the MIT Division of Comparative Medicine and MIT Committee on Animal Care. Female NOD/SCID/IL2R^null^ (NSG) mice were purchased from Jackson Laboratory and housed in pathogen-free conditions within MIT animal facilities. All mice were 8–12 weeks old upon initial experimental use. When initial tumor burden was detectable, mice were randomized into experimental groups to approximately equalize the starting means and standard deviations across groups.

### Primary human cells

All studies conducted with human tissue followed MIT Committee on the Use of Humans as Experimental Subjects (COUHES) policies. Peripheral blood mononuclear cells (PBMCs) from de-identified healthy donors were purified from leukapheresis products purchased from Stem Cell Technologies using EasySep Direct Human PBMC Isolation Kits (Stem Cell Technologies), following manufacturer instructions, prior to immediate cryopreservation. All primary cells were maintained in RPMI-1640 (ATCC) + 10% FBS + 100 U/mL P/S + 30 U/mL of recombinant human IL-2 (R&D Systems) at 37°C in 5% CO2.

### Library assembly

To generate scRNA-seq-compatible barcoded CAR constructs, randomized 8mer barcodes were inserted into the 5’UTR of an anti-CD19 backbone encoding a Flag epitope-tagged FMC63 scFv, IgG4 hinge, and CD28 transmembrane (TM) domain, along with an IRES-GFP element for the bicistronic expression of GFP, all within a pHIV lentiviral vector, as described previously.^26^ Downstream of the TM domain, BsmBI recognition sites flanking a LacZ domain were incorporated to allow for insertion of ICDs in subsequent cloning reactions. Barcodes were inserted via Gibson Assembly, after which the cloning product was electroporated into DH10B electrocompetent *E. coli* cells (ThermoFisher) to achieve high coverage of barcodes. The resulting barcoded pool was grown up, then used as a pre-barcoded destination vector for the construction of the caRNA-seq library. To create this defined library of ICD architectures, gene blocks encoding each ICD (codon optimized from the amino acid sequence corresponding to the intracellular portion of each respective immune receptor; see **Table S2**) were used as PCR templates for the addition of position-specific Golden Gate overhangs, allowing the single-step assembly of each ICD in a specified order into the pre-barcoded CAR backbone. Each variant was cloned in array and separately sequence verified.

### Lentivirus (LV) production

HEK293T cells grown to a confluence of 70-90% were transfected with a mix of pHIV CAR transfer plasmid, pMD2.g (VSV-G), and psPAX2 at a mass ratio of 5.6:1:3, complexed with polyetherimide (PEI) at a total DNA:PEI mass ratio of 1:3. Each CAR construct in the library was transfected separately in a 10cm dish, using 14 µg of transfer plasmid. Supernatant was collected at 48, 72, and 96 h post-transfection, after which collections from each time point were pooled, filtered through a 0.45 µm low protein-binding filter, and concentrated at 100,000*g* for 1.5 h. Concentrated LV was resuspended in serum-free media (OptiMEM, Gibco). LV titers were determined by transducing 1x10^5^ Jurkat cells with a dose series of concentrated LV in the presence of 8 µg/mL of dextran, then reading out GFP expression 48 h post-transduction.

### CAR T cell manufacturing

PBMCs were freshly thawed for each CAR T cell batch, after which bulk T cells T cells were enriched using EasySep Human T Cell Enrichment Kits (Stem Cell Technologies), then maintained in RPMI-1640 (ATCC) + 10% FBS + 100 U/mL P/S + 30 U/mL of recombinant human IL-2 (R&D Systems). For the pilot experiment in which CD4 cells with a defined CAR BC were spiked into CD8 cells (**Figure 1D-F**), CD4 and CD8 T cells were isolated separately using EasySep Human CD4+ or CD8+ Enrichment Kits (Stem Cell Technologies). Isolated T cells were immediately activated with DynaBeads Human T-Activator CD3/CD28 (ThermoFisher), using a 1:1 bead:cell ratio. One day after activation, concentrated LV encoding each CAR construct was added in the presence of 8 µg/mL of dextran, using a multiplicity of infection (MOI) of 5, based on functional titers calculated via titration on Jurkat T cells. Four days after initial activation, DynaBeads were removed from culture via magnetic separation, and CAR^+^ T cells were enriched via FACS (BD FACSAria II or Sony MA900) on the basis of GFP expression. Sorted cells were rested for 4 days following de-beading prior to use in any experiment (either direct characterization for tonic signaling studies, or antigen challenge). Throughout the manufacturing process, T cells were maintained at a density between 5x10^5^ and 2x10^6^ cells/mL.

### In vitro rechallenge assays

CAR T cells were cocultured with tumor cells (NALM6 or CD19^+^ A375) in IL-2-deficient media at an effector:target (E:T) ratio of 1:3, using 2.5x10^5^ CAR+ T cells and 7.5x10^5^ tumor cells in separate wells of a 24-well plate. Three days following the first antigen challenge, cells were sampled for flow cytometry such that CAR T cells and residual CD19^+^ tumor cells could be enumerated with counting beads (CountBright Plus Absolute Counting Beads, ThermoFisher). Based on these counts, another 2.5x10^5^ CAR+ T cells were resampled for a subsequent antigen challenge, again using 7.5x10^5^ total tumor cells (after accounting for any residual tumor burden). This process was repeated on day 5. On day 7, cells were collected, stained for CD19 expression and cell viability, and live GFP+ CAR T cells were sorted via FACS (Sony M900) for scRNA-seq. For arrayed NALM6 rechallenges, as shown in **Figure 2A**, this process was done separately for each library variant, and equal numbers of CAR^+^ cells were pooled on day 7 from each coculture prior to staining and sorting. For pooled A375 and NALM6 rechallenges, data from which is shown in **Figure 7E**, CAR T cells were pooled prior to the first antigen challenge.

### In vitro live killing assays

For NALM6 assays, flat-bottom 96-well plates were coated with 50 µL/well of 0.01% poly-L-ornithine (PLO) for 1 h at room temperature, after which the PLO was aspirated, and wells were dried for 0.5 h. 1x10^4^ mCherry^+^ NALM6 cells were seeded onto PLO-coated plates. For A375 assays, 1x10^4^ mCherry^+^ A375 cells were seeded 24 h prior to CAR-T addition on non-PLO-coated plates, over which we assumed the cell count to have doubled. CAR T cells were then added to tumor cells at an E:T of 1:1 or 1:10, and tumor cell growth was tracked longitudinally via red fluorescence on an Incucyte S3, with images taken every 3 h. Tumor burden was quantified as the total red area per well, after which values were normalized to initial total red area at the first time point. All killing assays were completed in IL-2 deficient media.

### In vitro cytokine secretion assays

Two days following antigen challenge as described above, supernatant was collected, cells were removed via centrifugation, and cytokine levels were measured using Human Uncoated ELISA Kits (ThermoFisher) targeting IFNγ, IL-2, or TNFα, following manufacturer instructions. Absorbance was read out on a Tecan Infinite M200 Pro plate reader, and concentrations of each cytokine was calculated based on standards included for each individual experiment.

### In vitro assessment of memory state and activation

Prior to antigen challenge (for analysis of tonic signaling), 24 h following antigen challenge, and 7 days after antigen challenge, cocultures were collected and stained for activation markers, CD69 and 4-1BB, as well as memory markers, CD62L and CD45RA. Expression levels were read out via flow cytometry.

### In vivo tumor models

For NALM6 leukemia studies, 8–12-week-old female NSG mice were injected via the tail vein with 5x10^5^ FFLuc^+^ NALM6 cells. Four days after tumor inoculation, mice were treated via tail vein injection with 2x10^5^ CAR^+^ cells, as determined by GFP expression, or 2x10^5^ mock-transduced T cells. Tumor burden was monitored longitudinally twice per week (or once per week if mice reached long-term remission) by intraperitoneal injection of IVISbrite D-Luciferin substrate (PerkinElmer) at a dose of 0.15 mg per gram of body weight, then measurement of bioluminescence with an IVIS Spectrum imaging system (PerkinElmer). Tumor burden was quantified by total photon counts, using the LivingImage software. Mice were monitored daily for signs of weight loss, discomfort, or morbidity, and euthanized when reaching humane endpoint, including weight loss greater than 20% of initial body weight.

For A375 melanoma studies, 8–12-week-old female NSG mice were inoculated subcutaneously in the right flank with 5x10^5^ CD19^+^ A375 cells. 21 days after tumor inoculation, mice were treated via tail vein injection with 5x10^5^ CAR^+^ cells, as determined by GFP expression, or 5x10^5^ mock-transduced T cells. Tumor burden was tracked longitudinally every 2-3 days via caliper measurement across two dimensions. Tumor area was approximated as A = L*W, while tumor volume was approximated as V = ½*L*W^2^, where L represents the longest dimension and W represents the shortest. Mice were euthanized when reaching humane endpoint, including weight loss as described above, signs of limited mobility, tumor ulceration, tumor area above 100 mm^2^, or tumor volume above 1000 mm^3^.

Tissues were harvested following humane euthanasia from the spleen of both NALM6- and A375- bearing mice, and the flank tumors of A375-bearing mice. Spleens were processed by mashing through a 70 µm cell strainer (ThermoFisher), followed by red blood cell lysis via treatment with ACK lysing buffer (ThermoFisher). A375 tumors were processed via an initial mechanical dissociation step using scissors, followed by enzymatic dissociation and tissue homogenization using a gentleMACS Human Tumor Dissociation Kit and Octo Dissociator (Miltenyi).

### Flow cytometry and cell sorting

Cells harvested from *in vitro* cultures or *in vivo* tissues, as described above, were washed in PBS before an initial round of staining with Live/Dead Fixable viability dyes (ThermoFisher) in PBS, for 15 min at 4°C. Surface markers were subsequently stained with antibody cocktails diluted in FACS buffer, composed of PBS, 0.5% BSA, and 2 mM EDTA, for 20 min at 4°C. Stained cells were then washed twice with FACS buffer before analysis on a Cytoflex S (Beckman), or cell sorting on a BD FACSAria II or Sony MA900. For samples stained with CITE-seq antibodies, a similar process was followed using a pool of TotalSeq-C barcoded antibodies purchased from BioLegend included in the normal fluorescent antibody mix. Following staining and washing, cells were sorted as described prior to scRNA-seq.

### Amplicon sequencing of CAR BCs

Genomic DNA from CAR T cells harvested directly from *in vitro* cultures or *in vivo* tissues was isolated using a PureLink Genomic DNA Mini Kit (ThermoFisher), following manufacturer’s instructions. CAR BCs were amplified using primers specific for the 5’UTR within the CAR constructs, which also included handles with Illumina-compatible adapters. PCR reactions were purified using AMPure XP beads (Beckman), following manufacturer’s instructions, short-read sequencing via the Genewiz Amplicon-EZ service.

### scRNA-seq: Cell encapsulation and library preparation

Following FACS enrichment, purified CAR T cells were counted, and approximately 10,000 cells per lane were encapsulated using a Chromium Next GEM Single Cell 5’ Kit v2, loaded onto a Chromium X (10X Genomics). To maximize capture of CAR BCs, a custom reverse transcription (RT) primer specific for the CAR transcript (targeted to the scFv, such that it would bind all ICD variants; 5’-AAGCAGTGGTATCAACGCAGAGTACCAACCACTCAAGTCCCTTTC-3’) was spiked into the RT reaction mix at a final concentration of 0.1 µM. Aside from this exception, the gene expression (GEX) and feature barcoding (FB; for samples stained with CITE-seq antibodies) libraries were generated according to kit instructions.

To generate CAR BC libraries, we used a nested PCR strategy similar to that used for enrichment of TCR transcripts in the generation of scRNA-seq VDJ libraries. In brief, the fraction of cDNA containing full-length transcripts was input into an initial round of PCR, with a universal forward primer (5’-GATCTACACTCTTTCCCTACACGACGC-3’) that binds upstream of the cell barcode on all transcripts and an “outer” CAR-specific reverse primer (5’- GCGCTAAGACTGGAAGTTGTTTGC-3’). This PCR product was cleaned using a 0.5/1X SPRI selection. This was then input into a second PCR reaction to further enrich for CAR BCs, using the same universal forward primer and an “inner” CAR-specific reverse primer upstream of the original which also includes Illumina adapters (5’- GTGACTGGAGTTCAGACGTGTGCTCTTCCGATCTGCAAAGCCATGGTGGCTCTAGA-3’), for downstream indexing and sequencing. The second PCR product was purified by a left-sided 1.2X SPRI selection. Sample indexing can then be performed following the same protocol as the 10X gene expression libraries, doing a final clean up with a left-sided 1.2X SPRI selection.

The resulting GEX, FB, and CAR BC libraries were pooled to achieve approximately 30,000, 5,000, and 1,000 reads per cell, respectively. Exact ratios depended on the number of cells encapsulated and the exact flow cell used. Final Illumina pools were sequenced on a NovaSeq6000 (Illumina).

### scRNA-seq analysis

Raw sequencing reads were aligned to the Genome Reference Consortium Human Build 38 (GRCh38) and a cell-feature matrix was constructed using CellRanger software (v6.1.2–7.0.1). All downstream analysis used the Seurat package (v4.3.0.1).^57^ Initial filtering of low-quality cells was done on the basis of mitochondrial gene content and number of unique genes detected, with thresholds set in a dataset-specific manner based on distributions of each metric; ultimately, cells with greater than 5–10% of reads mapped to mitochondrial genes or less than 1000–2500 unique genes were filtered out. For cells stained with a sample-specific hashing antibody, samples were demultiplexed based on antibody hash reads using the HTODemux algorithm, as described previously^58^ and implemented in Seurat. Lastly, cells were assigned to specific CAR variants on the basis of CAR BC reads using the MULTIseqDemux algorithm with auto-thresholding, as described previously^59^ and implemented in Seurat. For all datasets described in this study, only cells assigned to a single CAR by the MULTIseqDemux algorithm were analyzed, with any cells assigned as negatives or multiplets filtered out.

GEX data was normalized and scaled separately for each dataset using the Seurat SCTransform function. To ensure cells did not cluster based on T cell clonotype, all TCR-related genes were filtered out of the VariableFeatures identified by the SCTransform function. Principal component analysis (PCA) was then performed on the scaled data, using only the remaining VariableFeatures. To address batch effects between donors within our arrayed NALM6 rechallenge datasets, we utilized canonical correlation analysis (CCA) integration as implemented in the Seurat package;^57^ all other datasets in which only one donor was available were processed separately and thus did not require batch correction. To identify different transcriptional states, cells were then clustered using the Leiden algorithm on the basis of the first 30 PCs, utilizing weighted nearest neighbors clustering to incorporate CITE-seq data, when available, and shared nearest neighbors clustering otherwise. Enrichment of cells expressing each CAR variant within the resulting clusters was calculated via chi-squared test. Differentially expressed genes between different cell clusters, as well as different groups (such as CAR variants or CAR classes), were identified by Wilcoxon rank-sum test comparing all cells in each population of interest, as implemented in the FindMarkers Seurat function. Pathway enrichment analysis was completed on lists of differentially expressed genes using gene set enrichment analysis (GSEA), running pre-ranked analyses with ranked gene lists weighted by the Bonferroni-adjusted *P* value, testing against canonical pathways, as described previously.^60^ To score cells on the activity of pre-defined gene sets, the UCell package was used;^56^ the specific gene sets utilized in this study are described in **Table S5**.

To process the pooled NALM6 rechallenge dataset in the context of the associated arrayed reference, as depicted in **Figure S2B-D**, we utilized Seurat’s projection pipeline.^61^ In brief, we clustered the arrayed NALM6 reference as described above, using only Donor 2 cells, as this was the matched donor for the pooled dataset. Following normalization and scaling of the pooled dataset as described above, each cell in the pooled context was mapped to the arrayed UMAP space, using the UMAP model computed for the arrayed dataset as input into the MapQuery function. This function calculates scores for each cell in the query dataset based on the confidence of the mapping to the reference space; we filtered out cells from the pooled dataset with confidence scores below 0.75.

### GRN definition and delineation into CAR classes

GRNs were identified using the SCENIC pipeline, as described previously.^33^ In brief, raw UMI counts for all protein coding genes were exported for all cells in the NALM6 *in vitro* rechallenge dataset, following the QC described above. Genes were further filtered out of the input dataset if they were detected in less than 1/1000 cells. This processed matrix was input into the SCENIC algorithm, along with a list of curated human TFs (downloaded from the SCENIC documentation), to identify coexpressed regulons using the GRNBoost2 algorithm. Target genes within regulons were then pruned based on the presence of sequence motifs for each TF, using RcisTarget. Given the stochastic nature of these algorithms, we repeated this process over 100 iterations; to converge on a final set of TFs and targets, we discarded all TFs detected in <80% of iterations, then filtered out all targets for each TF that were detected in <50% of iterations. This resulted in a final set of consensus regulons, consisting of TFs and their associated target genes (provided in **Table S4**). Each cell was then scored on these consensus regulons, using either the AUCell algorithm, as implemented in the SCENIC package, or the UCell package, which is thought to be highly correlated to AUCell;^56^ these regulon scores were taken as representative of GRN activity within each cell. CARs were then grouped into classes on the basis of these GRN scores, where the average regulon score was calculated for each CAR, scaled, and k-means clustered using *k =* 7. Only regulons detected in more than 100 cells were used for k-means clustering (121 of 123 regulons).

### Pseudobulk analysis

To compute pseudobulk transcriptional profiles across CARs or CAR classes, raw UMI counts for all VariableFeatures, determined as described above, were aggregated across all cells within each population. These aggregated counts were input into the DESeq2 package for normalization and transformation via the *rlog* function. These transformed values were used as input into PCA, as well as to calculate pairwise Euclidean distances and Pearson correlations between groups. To calculate the tonic signaling score introduced in this study, we used the Euclidean distance between each pseudobulked CAR and the non-signaling negative control CAR, CAR1, within the resting unstimulated dataset. To compute pseudobulk GRN activity across CARs or CAR classes, GRN activity scores, calculated as described above, were averaged across all cells within each population. These average data points were then Z-scored for input into PCA, or to calculate pairwise Euclidean distances and Pearson correlations, as described above.

### Partial least squares regression (PLSR)

To identify transcription factor activity informative of phenotype, we performed partial least squares regression (PLSR). In particular, we regressed survival time (dependent variable) on SCENIC regulon activity scores (multivariate independent variables). While the dependent variable was measured from biological replicates for each candidate CAR, the independent variable was inferred from transcriptomic data for all CARs at single-cell resolution. Thus, to set up the model, we first aggregated survival time across replicates by the median. Next, we pseudo- bulked activity scores across CARs by the average. Finally, we took the median survival time for a candidate CAR as representative of the entire CAR class, allowing us to assign each pseudo- bulked CAR a corresponding survival time according to its CAR class membership. In total, this resulted in 35 observations across all phenotype – CAR combinations.

Next, prior to PLSR, we performed feature selection using least absolute shrinkage and selection operator (LASSO) regression with the *glmnet* (v4.1-8) package in *R* (v4.3.2). We retained only those features that were consistently selected (i.e., non-zero) across 1000 iterations. In each iteration, we performed leave-one-out cross-validation to identify the optimal regularization penalty value according to that which maximized the average cross-validated error. LASSO feature selection identified 14 regulons out of 121 total.

Finally, we ran PLSR with these retained features using the *ropls* (v1.34.0) package. Components were added until the dependent variable’s coefficient of determination (R^2^Y) < 0.01 or the cross-validated coefficient of determination (Q^2^Y, leave-one-out cross-validation) < 0.05. This identified two components. The model significance was assessed by comparing the model’s R^2^Y and Q^2^Y against a null distribution generated by 1000 permutations shuffling the response variable. Additionally, the RMSE was compared against a null distribution generated by 1000 iterations of randomly selected features from the original 121. All code associated with these analyses are available upon request.

## RESOURCE AVAILABILITY

### Lead contact

Further information and requests for resources and reagents should be directed to and will be fulfilled by the lead contact, Michael Birnbaum.

### Materials availability

Plasmids generated in this study are available upon request.

### Data and code availability

scRNA-seq data have been deposited at GEO (GSE264681) and are publicly available as of the date of publication. NGS data measuring CAR BC frequencies is deposited under the accession number PRJNA1102443. Code used to process scRNA-seq data is deposited to GitHub, available at https://github.com/birnbaumlab/caRNA-seq. Any additional information required to reanalyze the data reported in this paper is available from the lead contact upon request.

## QUANTIFICATION AND STATISTICAL ANALYSIS

Statistical details for all experiments can be found in the figure legends. Ns = not significant, *<0.05, **<0.01, ***<0.001, ****<0.0001. Graphical schematics and data figures were drawn and compiled using Adobe Illustrator.

### Supplemental information

Figures S1-S6 and Table S1-S3.

Table S4-S7 included as separate Excel files.

## Notes

### Competing Interest Statement

M.E.B. is a founder, consultant, and equity holder of Kelonia Therapeutics and Abata Therapeutics. K.S.G. is currently employed at Ginkgo Bioworks, Inc. M.V.M is an inventor on patents related to adoptive cell therapies, held by Massachusetts General Hospital (some licensed to Promab and Luminary) and University of Pennsylvania (some licensed to Novartis).
MVM receives Grant/Research support from Kite Pharma and Moderna, serves as a consultant for multiple companies involved in cell therapies, holds equity in 2SeventyBio, A2Bio, Affyimmune, BendBio, Cargo, Century Therapeutics, Neximmune, Oncternal, and TCR2 and serves on the Board of Directors of 2Seventy Bio. The other authors declare no competing interests.

