## Supplemental Information for "Library-based single-cell analysis of CAR signaling reveals drivers of *in vivo* persistence"

Figures S1-S6 and Tables S1-S3.
Tables S4-S7 included as separate Excel files.

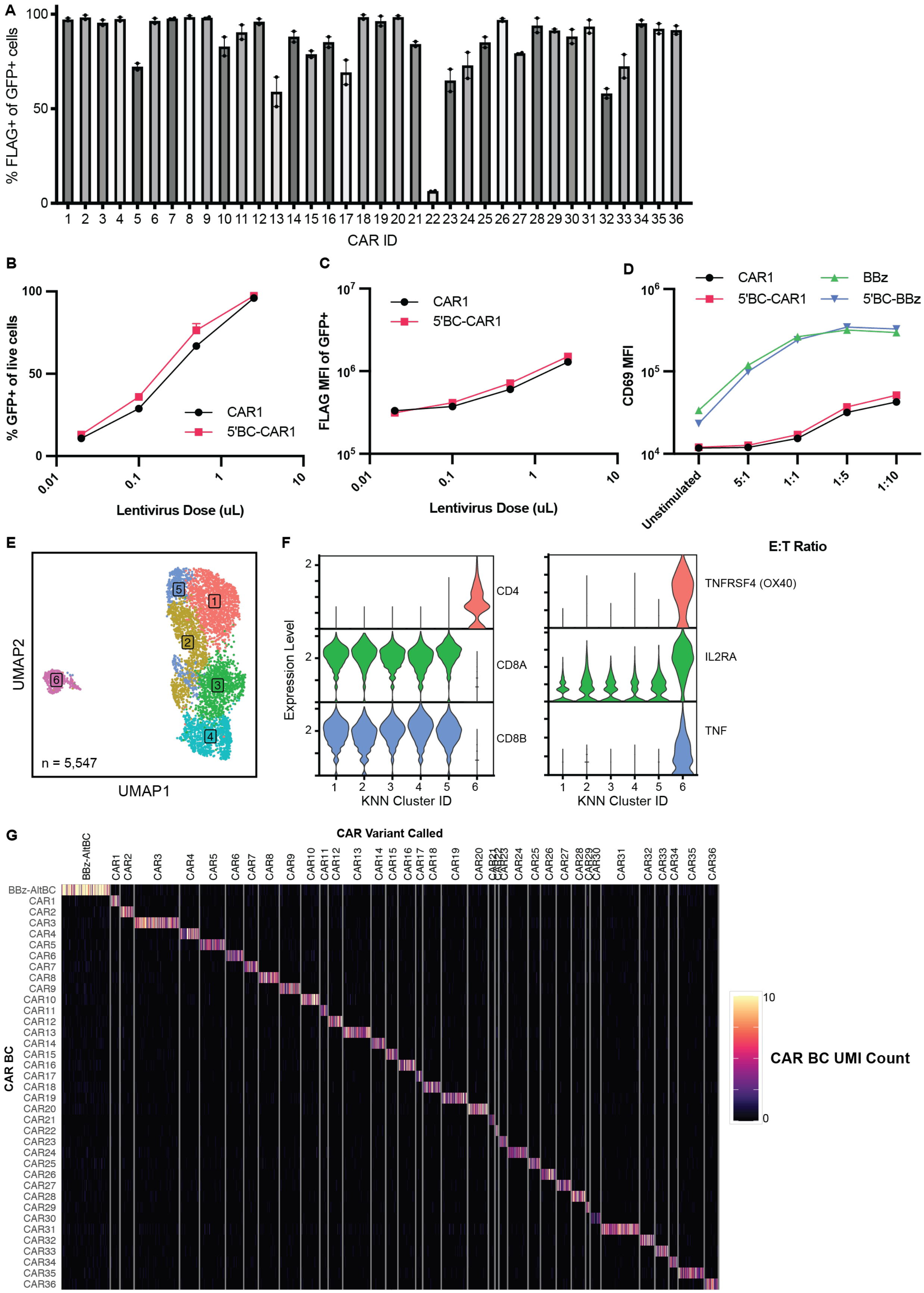

A
% FLAG+ of GFP+ cells
CAR ID
B
% GFP+ of live cells
Lentivirus Dose (uL)
CAR1
5'BC-CAR1
C
FLAG MFI of GFP+
Lentivirus Dose (uL)
D
CD69 MFI
CAR1
5'BC-CAR1
BBz
5'BC-BBz
Unstimulated
5:1
1:1
1:5
1:10
E:T Ratio
E
UMAP2
UMAP1
n = 5,547
F
Expression Level
CD4
CD8A
CD8B
TNFRSF4 (OX40)
IL2RA
TNF
KNN Cluster ID
G
CAR Variant Called
CAR BC
CAR BC UMI Count

**Figure S1. Characterization and validation of the caRNA-seq platform, related to Figure 1.**
(A) Percentage of Jurkat T cells staining positive for FLAG epitope tag incorporated into each construct, by flow cytometry. Datapoints represent two different lentiviral batches and independent transductions.
(B) Expression of transduction marker (GFP) on Jurkat T cells transduced with increasing doses of lentiviruses encoding an unbarcoded CAR1 or CAR1 with a library of random barcodes in the 5'UTR.
(C) FLAG mean fluorescence intensity (MFI), indicative of CAR surface expression, of GFP+ cells in (B).
(D) Expression of the activation marker CD69, quantified by MFI, in Jurkat T cells expressing 5'UTR barcoded and unbarcoded versions of CAR1 (non-signaling negative control) and 41BB-CD3z CAR (signaling competent positive control, BBz) following 24 h challenge with CD19+ NALM6 leukemia cells at various effector-to-target (E:T) ratios.
(E-G) Extended results of CD4 spike-in experiment. UMAP embeddings of all cells called to a single CAR variant (n = 5,547 cells) are presented in (E). Cells are colored by transcriptional state, as determined by unsupervised clustering (K-Nearest Neighbors, KNN) of gene expression. (F) Violin plots depicting expression of markers overexpressed and underexpressed in the activated CD4 cluster 6, including CD4, CD8, as well as effector-related molecules. (G) Heatmap of CAR BC expression across all 37 different CAR BCs (36 CARs in the library, as well as alternately barcoded BBz, BBz-AltBC), with cells grouped based on the CAR variant assigned. Columns represent individual cells, with each unique CAR BC on a separate row.
All datapoints in (A-D) are presented as mean ± SEM, representative of n = 3 technical replicates.

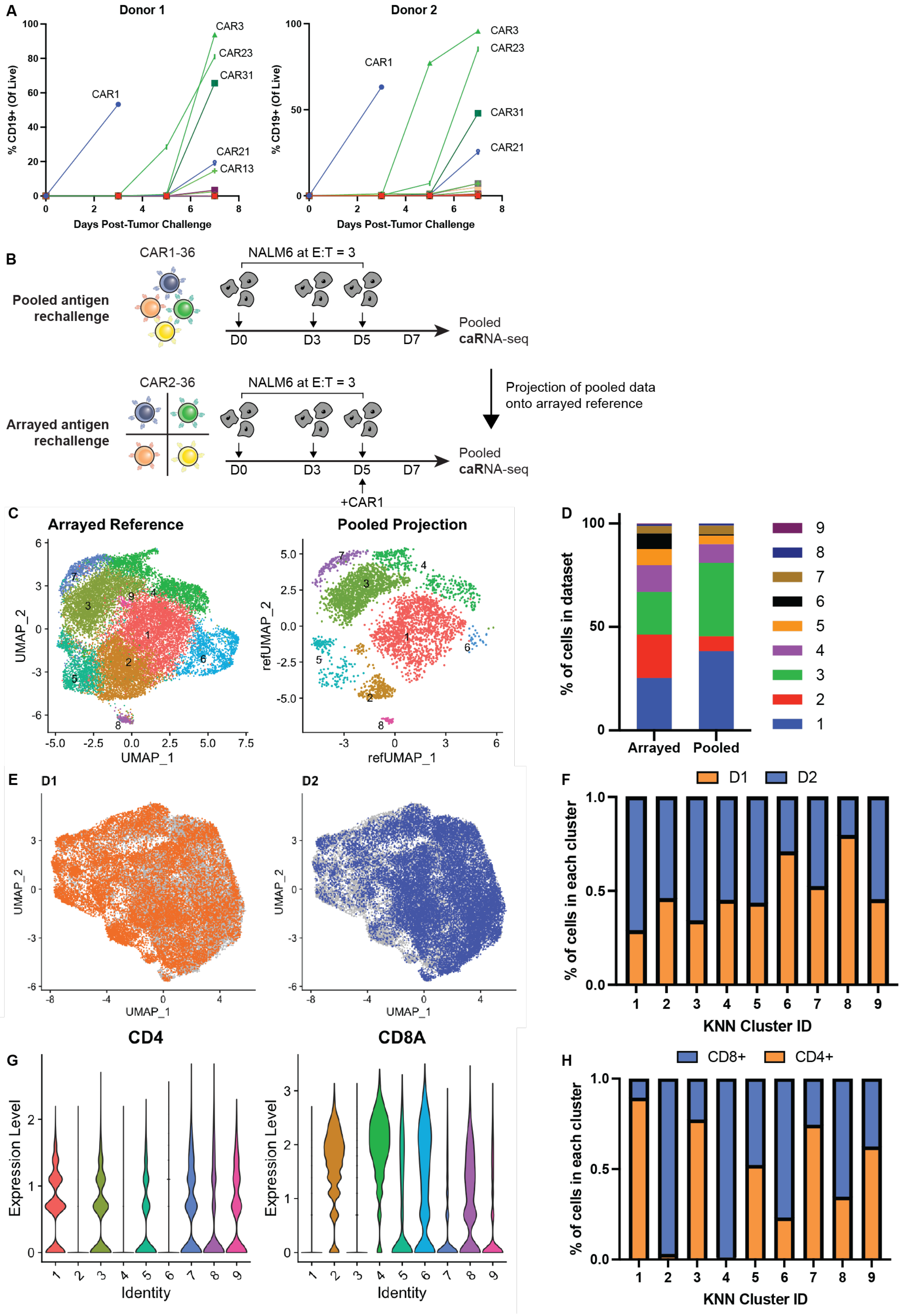
A
Donor 1
Donor 2
% CD19+ (Of Live)
Days Post-Tumor Challenge
CAR1
CAR3
CAR23
CAR31
CAR21
CAR13
B
CAR1-36
NALM6 at E:T = 3
Pooled antigen rechallenge
D0
D3
D5
D7
Pooled caRNA-seq
CAR2-36
Arrayed antigen rechallenge
+CAR1
Projection of pooled data onto arrayed reference
C
Arrayed Reference
Pooled Projection
UMAP_1
UMAP_2
refUMAP_1
refUMAP_2
D
% of cells in dataset
Arrayed
Pooled
E
D1
D2
F
% of cells in each cluster
KNN Cluster ID
G
CD4
CD8A
Expression Level
Identity
H
CD8+
CD4+

**Figure S2. Establishment of *in vitro* rechallenge assay to characterize ICD-intrinsic transcriptional programs, related to Figure 2.**

(A) The percentage of live $CD19^+$ tumor cells detected within each CAR variant's respective coculture, tracked over the course of the one-week NALM6 rechallenge for each donor. Only CARs with over 10% $CD19^+$ cells detected are labeled.

(B-D) Comparison of arrayed and pooled coculture reveals pooled noise. Experimental schematic depicting our comparison of pooled and arrayed NALM6 rechallenges, including projection of the pooled dataset onto the arrayed reference using the Seurat package's "projection" algorithm.[61] (C) Projection results showing the UMAP space defined by the arrayed reference (n = 19,020), and the pooled cells projected into this reference space (n = 4,210). Cells from the arrayed dataset are colored by transcriptional state, and cells from the pooled dataset are colored by inferred transcriptional state following projection. Only cells from the same donor (D2) were used to construct the arrayed reference. Cells from the pooled dataset with a projection confidence score lower than 0.75 were filtered out. (D) Percentage of cells assigned to each transcriptional cluster in the arrayed reference and the pooled projection. Cells from the pooled dataset with a projection confidence score lower than 0.75 were filtered out.

(E-H) Distribution of donor and CD4/8 subsets across arrayed rechallenged dataset. (E) Donor of origin overlaid on each cell in the UMAP space following batch correction and integration. (F) Percentage of cells in each transcriptional cluster from each donor. (G) Violin plots depicting CD4 and CD8 gene expression across different transcriptional clusters. (H) Percentage of cells in each cluster assigned as CD4+ and CD8+ based on expression of CD4, CD8A, and CD8B genes.

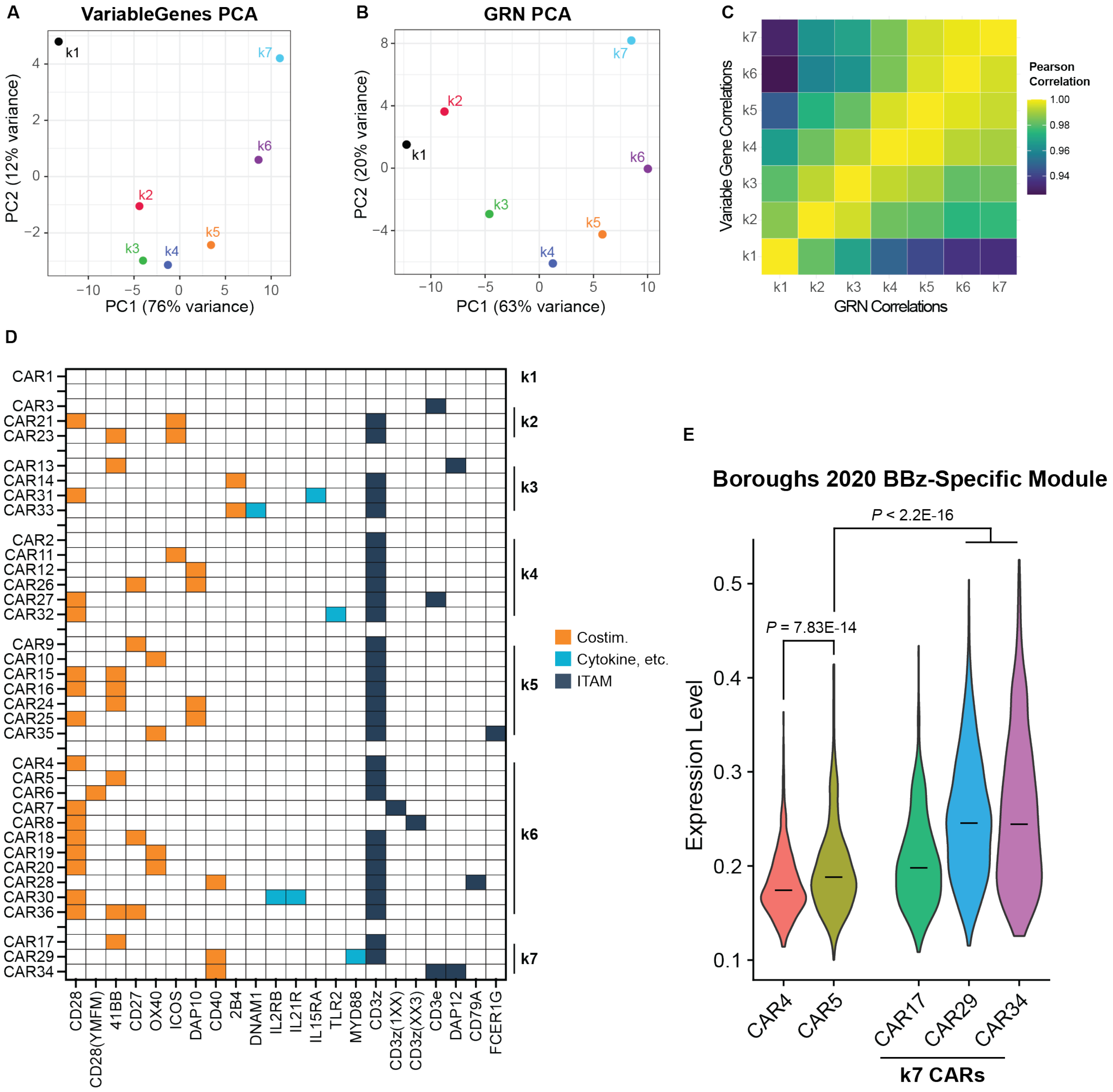


**Figure S3. High-level comparisons between CAR classes, related to Figure 3.**
(A-B) Principal component analysis (PCA) representing pseudobulked gene expression profiles averaged across all cells in each CAR class defined in Figure 3B, using (A) the expression of all variable genes or (B) SCENIC AUC scores representing gene regulatory network (GRN) activity.
(C) Pairwise Pearson correlation values comparing each CAR class with the others. Correlations were calculated using all variable genes (upper triangle), or GRN scores (lower triangle).
(D) ICD compositions grouped into the different CAR classes identified in **Figure 3B.**
(E) Violin plots depicting expression of a gene set shown to differentiate 28z and BBz CARs[10] across cells expressing 28z (CAR4), BBz (CAR5), and k7 CARs (CAR17, 29, and 34). Cells were scored using the *UCell* algorithm.[56] Bars represent median scores across all cells expressing the respective CAR. *P* values were calculated by Wilcoxon rank-sum test.

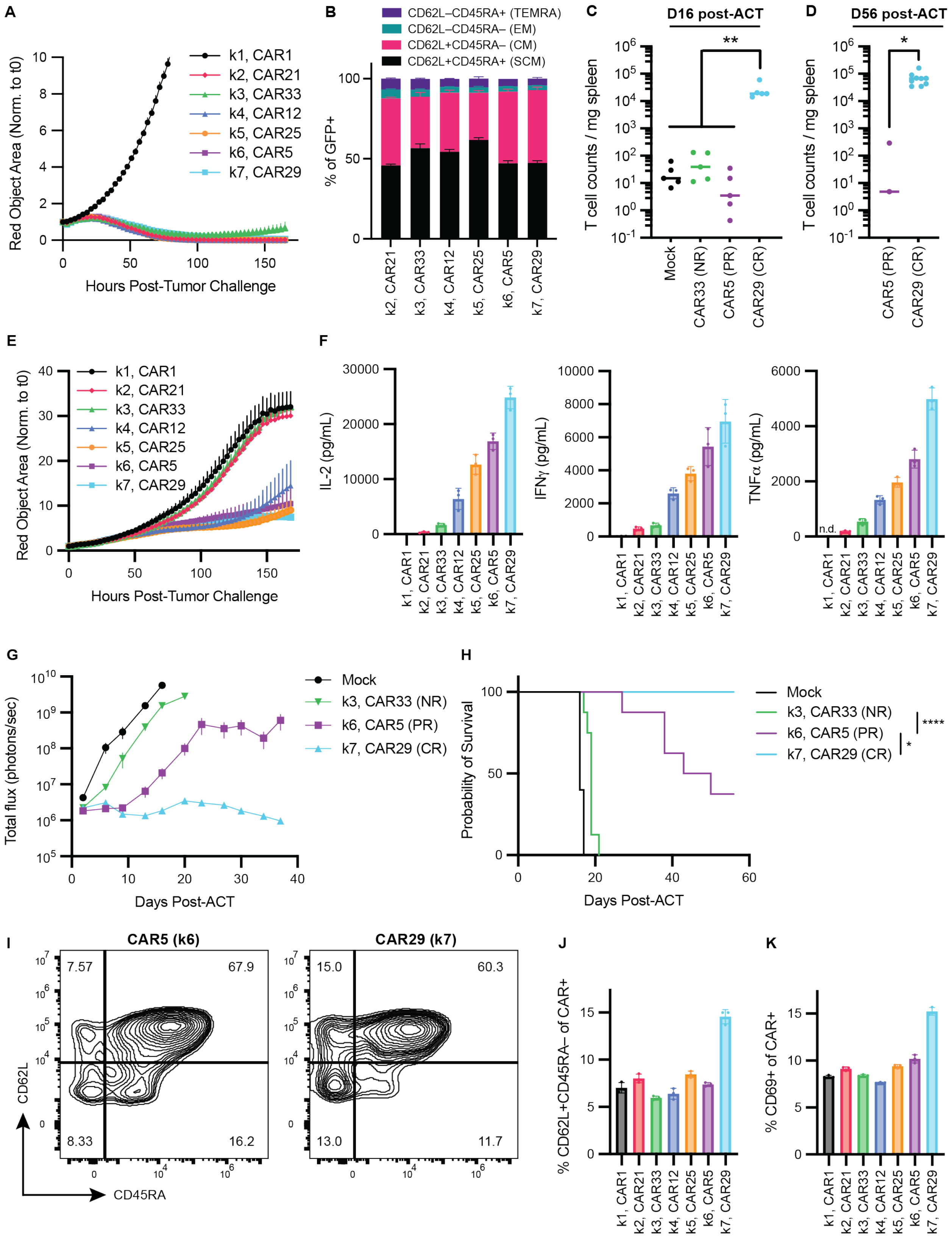


**Figure S4. Extended functional characterization of CAR candidates from each CAR class, related to Figure 4.**

(A) Longitudinal control of NALM6 tumor burden at an effector-to-target (E:T) ratio of 1:1 over seven days, as measured by live cell imaging, across candidate CARs from each CAR class. Data is shown as mean ± SEM, representative of n = 3 technical replicates, from a representative donor.
(B) Expression of surface markers indicative of memory state seven days after high tumor burden NALM6 challenge (effector-to-target, E:T = 1:10) across candidate CARs from each CAR class. CD62L and CD45RA expression was measured by flow cytometry to classify CAR T cells into stem cell memory (SCM), central memory (CM), effector memory (EM), TEMRA subsets. Data is shown as mean ± SEM, representative of n = 3 technical replicates, from a representative donor.
(C) T cell abundances at day 16 post-treatment, measured for mice treated with $2x10^5$ CAR T cells expressing CAR33, CAR5, CAR29, or mock-transduced. In terms of efficacy, each of the CARs elicits no response (NR), partial response (PR), or complete response (CR), respectively. Data shows median ± SEM, aggregated across n = 5 mice per group.
(D) Same data as in (C) but at day 56 post-treatment, at which point only CAR29-treated mice (*n* = 9 mice) and a small set of CAR5-treated mice (*n* = 3) had survived in this cohort.
(E-K) Functional characterization of candidate CARs in an alternate donor. (E-F) Functional characterization of candidate CARs from each CAR class identified in **Figure 3**, following NALM6 tumor challenge (E:T = 1:10). Longitudinal control of tumor burden over seven days, as measured by live cell imaging, is shown in (E), and cytokine secretion levels at 48h post-stimulation, as measured by ELISA, are shown in (F). (G-H) Characterization of in vivo efficacy driven by three candidate CARs, one that was characterized as a non-responder (NR, CAR33), partial responder (PR, CAR5), and complete responder (CR, CAR29) (n = 8-9 mice per group). Mice bearing NALM6 tumors were treated at a dose sub-curative for a 19BBz/CAR5 control CAR, and tumor burden was tracked longitudinally by bioluminescence. Tumor growth curves are shown in (G), while overall survival is shown in (H). P values were calculated by log-rank Mantel-Cox test, prior to Bonferroni multiple hypothesis correction using all possible pairwise comparisons (*P < 0.05; **P < 0.01; ****P < 0.0001) (I-K) Functional characterization of candidate CARs at rest. Representative flow plots showing expression of memory surface markers for k6 and k7 candidates in the absence of antigen are shown in (I) and quantified across all CARs in (J) as the percentage of CD62L+CD45RA– CARs. Expression of the activation marker CD69 at rest is quantified in (K). P values were determined by one-way ANOVA with Tukey's multiple comparisons test (****P < 0.0001).
All pooled data is shown as mean ± SEM. In vitro studies in (A-B, E-F, and I-K) show three technical replicates from a representative donor.

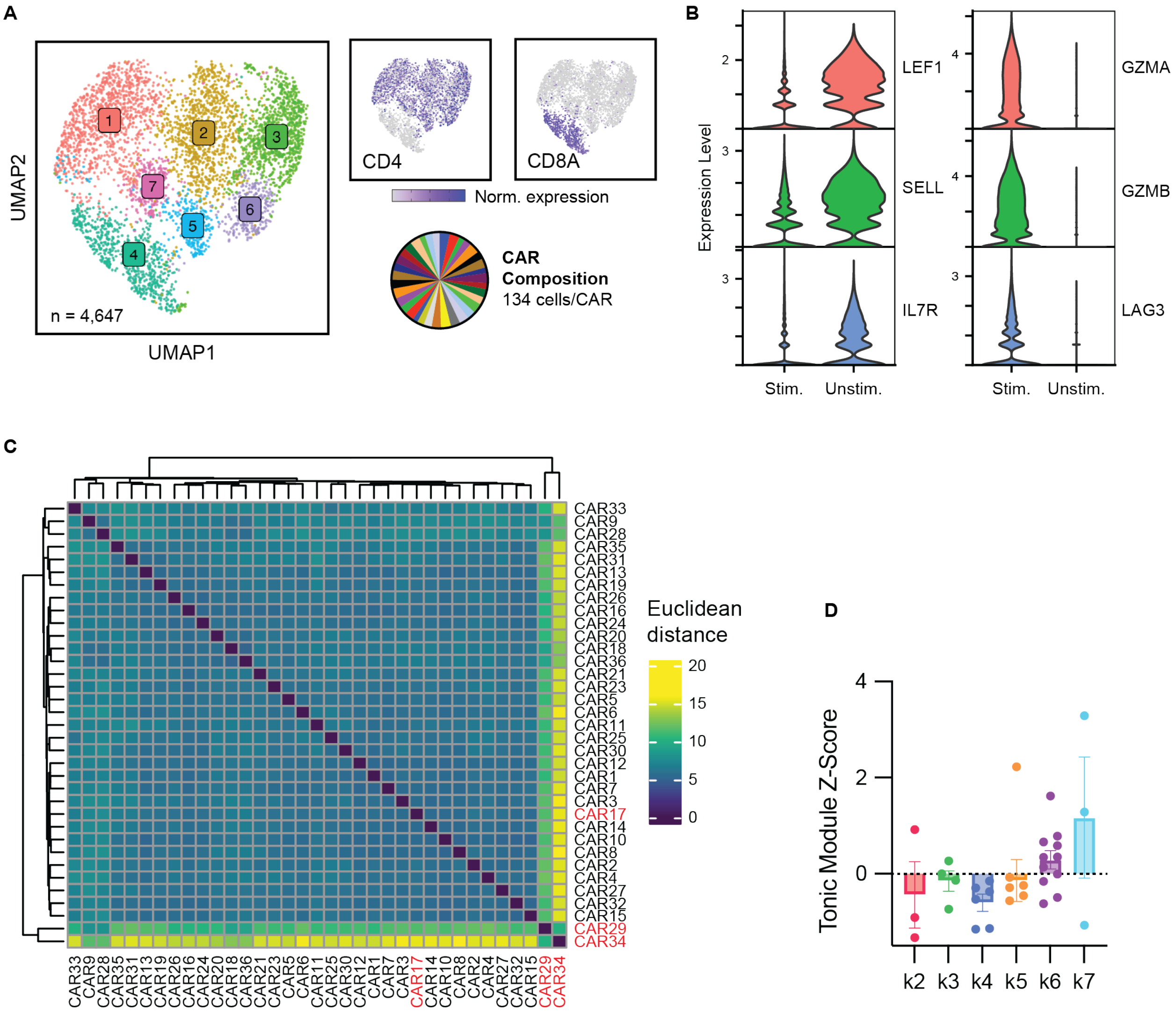


**Figure S5. Transcriptional characterization of CAR library at rest, related to Figure 5.**

(A) UMAP embeddings of all cells confidently called to a single CAR variant (n = 4,647 cells) in the resting dataset. Cells are colored by cell state (left), as determined by unsupervised clustering (K-Nearest Neighbors, KNN), and CD4 and CD8A RNA expression are shown overlaid (right). Proportions of cells assigned to each CAR is shown as a pie chart.

(B) Violin plots depicting normalized expression of genes related to T cell memory and stemness (left), as well as effector T cell molecules (right), in the antigen rechallenged dataset (Stim.) and the resting dataset (Unstim.).

(C) Pairwise Euclidean distances computed between each CAR in the library using by-CAR pseudobulked average expression profiles of all variable genes. The distances from CAR1 in this heatmap are taken as a tonic signaling score, and presented in Figure 5D.

(D) Geneset *z*-scores representing expression of a CD3z-mediated tonic signaling signature[10] across each CAR class at rest. Each datapoint represents a single pseudobulked CAR within each class, as calculated by the *UCell* algorithm.[56]

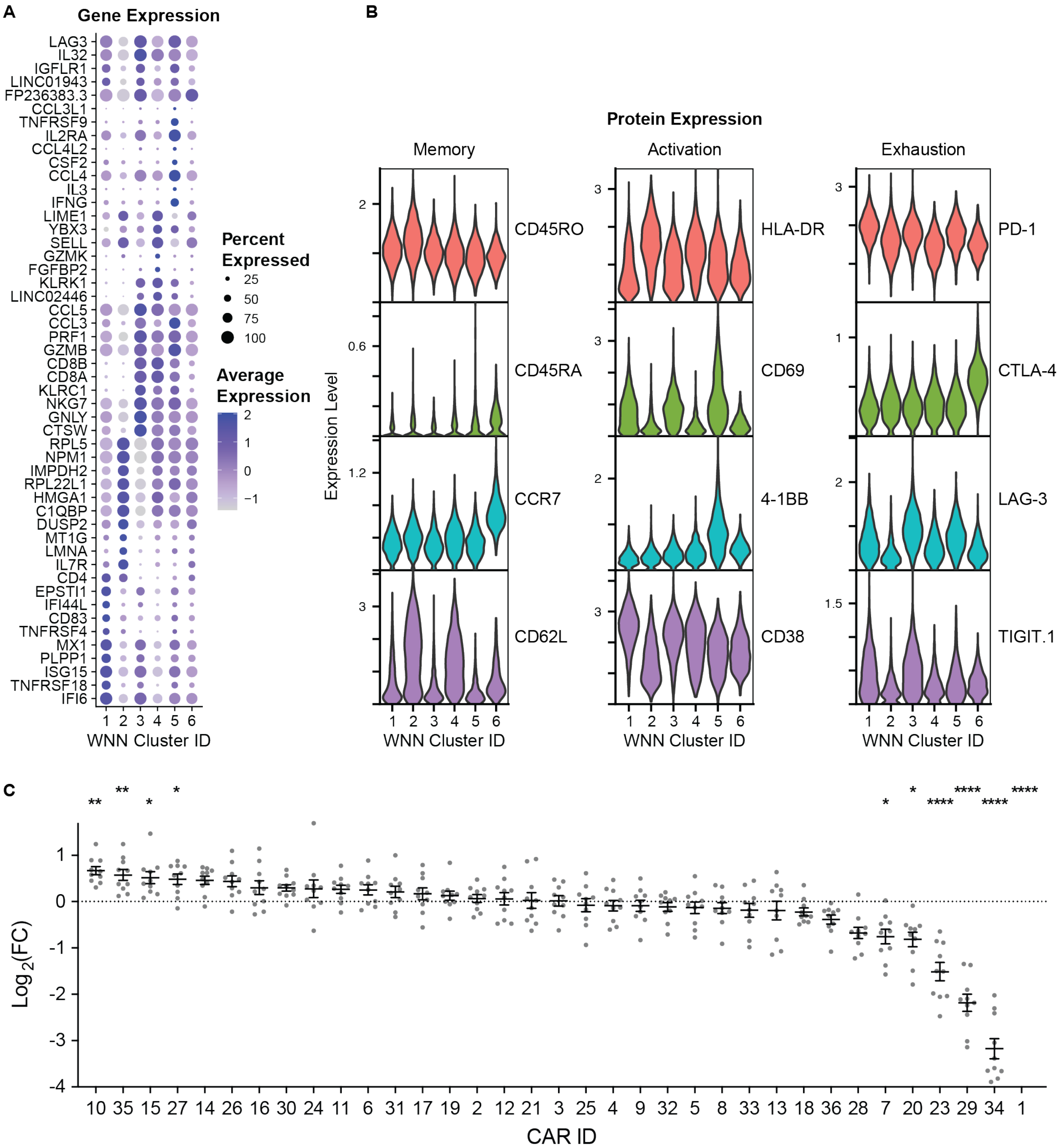


**Figure S6. Extended transcriptional characterization of caRNA-seq library in the in vivo solid tumor microenvironment, related to Figure 7.**

(A) Dot plot depicting average levels of gene expression for the top 10 differentially expressed genes in each cluster identified in Figure 7C. Color intensity represents average level of expression within each cluster, while size of the dot represents the percentage of cells in each cluster with detected expression of the gene.

(B) Protein expression levels, as detected by CITE-seq antibody staining, across clusters, depicted in violin plots. Markers are grouped into memory, activation, and exhaustion.

(C) Enrichment of CAR variants within the solid tumor. Data are represented as fold change values from pre-injection proportions, as measured by next-generation sequencing (NGS) of CAR BCs, to those observed at tumor harvest. Data points represent different tumors from individual mice, compiling both NGS (n = 7 mice) and scRNA-seq cell proportions (n = 3 mice). P values were determined by one-way ANOVA with Tukey's multiple comparisons test (****$P < 0.0001$), where each CAR was compared to the enrichment observed in the CAR5/19BBz control.

### Table S1. Description of ICD architectures in the caRNA-seq library

| Name | ICD1 | ICD2 | ICD3 | ICD4 | Reference | Barcode | Notes |
|---|---|---|---|---|---|---|---|
| CAR1 | X | X | X | X | N/A | TCAGCTCG | Null CAR with no ICDs |
| CAR2 | CD3z | | | | N/A | GTTACCCC | 1st gen CAR control |
| CAR3 | CD3e | | | | N/A | AGGAAATC | 1st gen CAR control |
| CAR4 | CD28 | CD3z | | | N/A | ACCCACGT | FDA-approved (Yescarta) |
| CAR5 | 4-1BB | CD3z | | | N/A | AAATAACC | FDA-approved (Kymriah) |
| CAR6 | CD28(YMFM) | CD3z | | | Guedan 2020[15] | TACGTGAG | Grb2 motif mutated in CD28 |
| CAR7 | CD28 | CD3z(1XX) | | | Feucht 2019[16] | GAGACAGG | 2/3 ITAM-deficient CD3z |
| CAR8 | CD28 | CD3z(XX3) | | | Feucht 2019[16] | AAAGCACC | 2/3 ITAM-deficient CD3z |
| CAR9 | CD27 | CD3z | | | Song 2012[11] | AAGTAAGG | |
| CAR10 | OX40 | CD3z | | | Pulè 2005[12] | ACTCATTA | |
| CAR11 | ICOS | CD3z | | | Guedan 2018[13] | GGAGTGAC | |
| CAR12 | DAP10 | CD3z | | | Lynch 2017[21] | ACGACGAC | |
| CAR13 | 4-1BB | DAP12 | | | Ng 2021[22] | CTGACCTG | |
| CAR14 | 2B4 | CD3z | | | Xu 2019[23] | CCAGGCCA | |
| CAR15 | 4-1BB | CD28 | CD3z | | Zhao 2009[62] | TAAGAAAC | |
| CAR16 | CD28 | 4-1BB | CD3z | | Zhong 2010[63] | TACGTTCA | |
| CAR17 | 4-1BB | CD3z | CD3z | | Majzner 2020[64] | CATAAGAG | |
| CAR18 | CD28 | CD27 | CD3z | | Nair 2018[18] | AAAATACA | |
| CAR19 | CD28 | OX40 | CD3z | | Guercio 2021[65] | AACTATAA | |
| CAR20 | CD28 | CD3z | OX40 | | Hombach 2012[66] | GCAATTGT | |
| CAR21 | CD28 | ICOS | CD3z | | Nair 2018[18] | AAAATCAA | |
| CAR22 | ICOS | 4-1BB | CD3z | | Guedan 2018[13] | ATCTTACA | |
| CAR23 | 4-1BB | ICOS | CD3z | | Guedan 2018[13] | GCCAAGCA | |
| CAR24 | 4-1BB | CD3z | DAP10 | | Zhao 2019[67] | AAATATGC | |
| CAR25 | CD28 | CD3z | DAP10 | | Lv 2019[68] | GATCGCGC | |
| CAR26 | DAP10 | CD27 | CD3z | | Duong 2013[69] | TGATCTCG | |
| CAR27 | CD3e | CD28 | CD3z | | Wu 2020[14] | CCTTAATC | |
| CAR28 | CD79A | CD40 | CD3z | | Julamanee 2021[25] | AGTCATCT | |
| CAR29 | tMyD88 | CD40 | CD3z | | Prinzing 2020[19] | AAAGTATC | Truncated MYD88 domain |
| CAR30 | CD28 | tIL2RB | CD3z(YXXQ) | | Kagoya 2018[17] | CCGATAAA | Truncated IL2RB, STAT3 motif at C-terminus of CD3z |
| CAR31 | CD28 | IL15Ra | CD3z | | Nair 2018[18] | AGCATGTT | |
| CAR32 | CD28 | CD3z | tTLR2 | | Weng 2018[20] | GATACCCT | Truncated TLR2 domain |
| CAR33 | DNAM-1 | 2B4 | CD3z | | Huang 2020[24] | CTAAAAAT | |
| CAR34 | CD40 | CD3e ITAM | DAP12 | | Gordon 2022[26] | TCGGAGCA | |
| CAR35 | FCER1G | OX40 | CD3z ITAM3 | | Gordon 2022[26] | CTATAGTG | |
| CAR36 | CD28 | 4-1BB | CD27 | CD3z | Chang 2016[70] | GGATGAGA | |

### Table S2. Description of ICDs built into caRNA-seq library

| ICD | Uniprot accession number | Amino acid positions | Mutations to endogenous sequence | Notes |
|---|---|---|---|---|
| CD3z | P20963 | 52-164 | --- | |
| CD3z ITAM3 | P20963 | 131-159 | --- | |
| CD3z(1XX) | P20963 | 52-164 | Y111F, Y123F, Y142F, Y153F | |
| CD3z(XX3) | P20963 | 52-164 | Y72F, Y83F, Y111F, Y123F | |
| CD3z(YXXQ) | P20963 | 52-164 | L156Y, H157R, M158H | YXXQ domain (STAT3-binding motif) inserted into C terminal tail of CD3z |
| CD3e | P07766 | 153-207 | --- | |
| CD3e ITAM | P07766 | 178-205 | --- | |
| DAP12 | O43914 | 62-113 | --- | |
| CD79A | P11912 | 166-226 | --- | |
| FCER1G | P30273 | 45-86 | --- | |
| CD28 | P10747 | 180-220 | --- | |
| CD28(YMFM) | P10747 | 180-220 | N193F | Abrogates binding of Grb2 to YMNM motif |
| 4-1BB | Q07011 | 214-255 | --- | |
| CD27 | P26842 | 213-260 | --- | |
| OX40 | P43489 | 236-277 | --- | |
| ICOS | Q9Y6W8 | 162-199 | --- | |
| DAP10 | Q9UBK5 | 70-93 | --- | |
| CD40 | P25942 | 216-277 | --- | |
| 2B4 | Q9BZW8 | 251-370 | --- | |
| DNAM1 | Q15762 | 276-336 | --- | |
| tIL2RB | P14784 | 266-337 and 529-551 | --- | Truncated via internal deletion, leaving JAK and STAT-binding sites |
| IL15RA | Q13261 | 229-267 | --- | |
| tTLR2 | O60603 | 626-784 | --- | Truncated, contains only TIR domain from TLR2 |
| tMYD88 | Q99836 | 1-158, 294-296 | --- | Truncated, TIR domain deleted |

### Table S3. Description of scRNA-seq datasets generated in this study.

| Dataset Name | Related Figure | Donor ID | Description |
|---|---|---|---|
| CD4 Spike-In | **Figure 1B-D** | 1 | Pilot study to assess CAR BC accuracy at library scale. Antigen-stimulated CD4 cells expressing an alternatively barcoded BBz (BBz-AltBC) were spiked into unstimulated CD8 cells expressing the full library of barcoded CAR variants at ~5%. |
| Arrayed NALM6 D1 | **Figure 2** and **Figure 3** | 1 | CAR library challenged with NALM6 *in vitro* in array on days 0, 3, and 5, then combined and sequenced in pool on day 7. |
| Arrayed NALM6 D2 | **Figure 2** and **Figure 3** | 2 | CAR library challenged with NALM6 *in vitro* in array on days 0, 3, and 5, then combined and sequenced in pool on day 7. |
| Resting | **Figure 5E-G** | 2 | CAR library sequenced at rest, in the absence of antigen. |
| Pooled NALM6 | **Figure 7E** and **Figure S5** | 2 | CAR library challenged with NALM6 *in vitro* in pool on days 0, 3, and 5, then sequenced on day 7. |
| Pooled A375 | **Figure 7E** | 2 | CAR library challenged with A375 *in vitro* in pool on days 0, 3, and 5, then sequenced on day 7. |
| In Vivo A375 | **Figure 7E-G** | 2 | CAR library harvested and sequenced from A375 flank tumors on day 10 post-treatment. Tumors from individual mice were stained with different hashing antibodies to allow deconvolution of biological replicates in sequencing space. Tumors were also stained with a panel of CITE-seq antibodies specific for T cell phenotypic markers. |

### Table S4. Gene regulatory networks identified via SCENIC in the arrayed NALM6 rechallenged dataset.

Provided as an attached Excel spreadsheet.

### Table S5. Literature-derived gene sets used in this study.

Provided as an attached Excel spreadsheet.

### Table S6. Differentially expressed genes between complete response (k7 CARs) and partial response (k5/k6 CARs) groups within the arrayed NALM6 rechallenged dataset.

Provided as an attached Excel spreadsheet.

### Table S7. Differentially expressed genes between responder (CAR5, CAR25) and non-responder (CAR12, CAR21, CAR29, CAR33) CARs within the *in vivo* A375 dataset.

Provided as an attached Excel spreadsheet.
